# Sorting States of Environmental DNA Reveals State-Specific Biodiversity Signals and Transport Patterns in Eight Alpine Watersheds

**DOI:** 10.64898/2025.12.17.694807

**Authors:** Anish Kirtane, Enrico van der Loo, Zora Doppmann, Kristy Deiner

## Abstract

Environmental DNA (eDNA) exists in three states: membrane-bound, adsorbed, and dissolved. These states differ in persistence and degradation, strongly influencing the interpretation of eDNA data. Despite this, they have rarely been separated and analyzed independently from environmental samples. We developed a state-sorting workflow to isolate and analyze them, applying it to samples from 221 sites from 58 streams across eight lake watersheds with COI and ITS metabarcoding to reveal differences in biodiversity content and transport dynamics.

Our results show that all three states contain both shared and unique taxonomic diversity for COI, and that plant DNA was only detected in the membrane-bound state. For COI, membrane-bound eDNA contained 87.8% of observed ASVs, far exceeding the adsorbed (37.2%) and dissolved (20.5%) states. Only membrane-bound eDNA showed evidence of downstream transport, but its extent varied among watersheds due to local hydrology. While upstream eDNA was transported to stream–lake confluences, lake surface samples showed marked turnover in community composition. Clarifying the fate of membrane-bound eDNA within lakes will enhance catchment-level detection from lake samples and understanding of lake hydrodynamics. Of the environmental parameters assessed, water temperature was most strongly aligned with changes in community composition between sites.

Most previous studies have likely captured the majority of the eDNA diversity within their samples by inadvertently targeting the membrane-bound state. This study demonstrates the utility of eDNA state-sorting, but the methods require further refinement. Analyzing states independently improves the interpretation of eDNA data and elucidates the processes governing eDNA fate and transport.

## Introduction

Assessing global biodiversity trends is challenging due to the vast landscapes supporting a wide range of biological diversity, each requiring specialized knowledge and survey methodologies that require substantial resources (Loh et al., 2005; Pereira & Cooper, 2006; Vié et al., 2009). Environmental DNA (eDNA) analysis enables taxonomic identification across the tree of life, often matching or surpassing the sensitivity of conventional methods cost-effectively to provide comprehensive biodiversity data (Deiner et al., 2021; Ruppert et al., 2019). It is hypothesized that as water flows through a landscape, it collects eDNA from terrestrial and aquatic sources (Deiner et al., 2016a; Jo & Yamanaka, 2022; Shogren et al., 2017). If eDNA is transported before it degrades, it may accumulate in predictable places (‘eDNA hotspots’) in the landscape. Lakes are strong contenders to be eDNA hotspots as multiple streams and rivers, carrying the eDNA collected across the landscape, drain into them. The large residence time of lakes may allow for the accumulation and homogenization of this eDNA information. However, empirical verification of this hypothesis and a comprehensive understanding of eDNA transport in lotic and lentic systems are a prerequisite for such eDNA-based landscape-level biodiversity surveys.

This transport of eDNA in lotic systems presents both opportunities and challenges for its use in biodiversity monitoring. Opportunities arise from the accumulation of eDNA from the catchment as water flows downstream. Streams can act as conveyor belts carrying the eDNA shed upstream to downstream sites, providing aggregated biodiversity information over a larger geographic area (Deiner et al., 2016a; Mächler et al., 2019; Pont et al., 2018; Wood et al., 2021). Thus, a sample collected downstream can offer biodiversity insights representing a broader area compared to an upstream sample, using the same resources (Deiner et al., 2016a). Conversely, a major challenge presented by eDNA transport is in disentangling the local occupancy of organisms (Burian et al., 2021). The challenge is to determine whether the captured eDNA was shed locally or transported to the sampling site, leading to potential misinterpretations of current local occupancy of organisms (Jo & Yamanaka, 2022; Wood et al., 2021). The heterogeneous release and downstream transport can lead to high eDNA turnover rates that can lead to false negative detections with modest sampling effort (Burian et al., 2021; Reji Chacko et al., 2023). Understanding the transport behavior of eDNA in streams is imperative to optimize the opportunities and limit the challenges.

Numerous studies have attempted to document the transport behavior of eDNA in lotic systems and reveal dramatically differing transport potentials. A meta-analysis estimated mean eDNA transport distances with a 95% confidence interval of 111.5–424.8 m, with most eDNA expected to travel <2 km under ordinary conditions, although extreme cases up to ∼13 km have been reported (Jo & Yamanaka, 2022). Hydrological modeling in rivers with larger discharge predicts potential eDNA transport distances of more than 100 km (Pont et al., 2018). The transport potential of eDNA is determined by its removal rate from the water column to concentrations below the limits of detection. This removal is facilitated via processes like settling, degradation, and dilution as water flows downstream which are influenced by hydrological parameters as well as the characteristics of eDNA itself (Barnes & Turner, 2016; Pont et al., 2018). Characteristics of the eDNA that impact its removal from the water column are its settling velocity and degradation rate (Pont, 2024; Pont et al., 2018), both of which are likely impacted by the state of eDNA (Jo & Yamanaka, 2022; Mauvisseau et al., 2022).

The behavior of eDNA is further complicated by the fact that eDNA is not a uniform entity; instead, it comprises multiple states—namely, membrane-bound, dissolved, and adsorbed (Mauvisseau et al., 2022; Nagler et al., 2022). Each state has distinct properties in terms of its degradation rates and settling velocity that can impact its transport potential (Jo & Yamanaka, 2022; Mauvisseau et al., 2022). The dissolved eDNA, for instance, appears to be highly vulnerable to degradation, whereas membrane-bound eDNA is shielded from extracellular degradative factors by cellular or organellar membranes reducing its degradation rate (Barnes & Turner, 2016; Mauvisseau et al., 2022; Nagler et al., 2022). Conversely, the adsorbed state in the right conditions can remain preserved over hundreds of years (Demaneche et al., 2001; Giguet-Covex et al., 2019; Paget et al., 1992). Membrane-bound DNA and adsorbed DNA can settle in non-turbulent conditions, unlike dissolved eDNA which tends to stay suspended (Barnes & Turner, 2016; Pont, 2024). However, it has not yet been empirically studied how estimates of richness and diversity are impacted by the state of eDNA captured, and how the transport of eDNA varies by its state. Methodologies developed previously have been tested and optimized for freshwater capture of different states (Kirtane et al., 2023; Kirtane & Deiner, 2024). While not completely isolating each state, these methods now enable the enrichment of eDNA states from the same water sample into their different states for independent analysis.

In this study, we sampled surface water from eight Swiss watersheds. For each watershed, the samples were taken from the headwaters to the lakes they drain into, resulting in a total of 221 sampling sites in 58 streams. We separated eDNA states from each water sample using optimized state-sorting methodologies and analyzed them with COI and ITS metabarcoding assays targeting metazoans and plants respectively. This approach allowed us to investigate the impact of the eDNA state on the diversity and richness of ASVs captured, and to understand how downstream transport is influenced by the eDNA state. Additionally, we tested whether surface water of lakes has eDNA from their inflowing streams and identified the states of eDNA being transported.

## Methods

### Field sampling and site selection

#### Choice of sampling sites

eDNA samples were collected from 221 sites in 58 streams in eight Swiss lake watersheds. Streams with little water flow or with a length from the headwaters to the lake below 2 km were excluded as they tend to be ephemeral in flow and would not summarize much biodiversity of a catchment. Starting at the highest accessible point, stream surface samples were collected along the stream at intervals of 1.5–3 km, extending downstream to the lake inlet. Inlet samples were collected from the stream 10–750 m upstream of the stream–lake confluence. The lake surface water samples were collected within 10 m from the stream-lake confluence. The exceptions were Lake Greifen and Hallwil, which do not permit boats closer than 50 m from the lake shore. Graphical illustrations of sampling locations as well as sample location maps can be found in Figure 1A and Figures S1- S8. All samples within each watershed were collected within one week.

**Figure 1:**
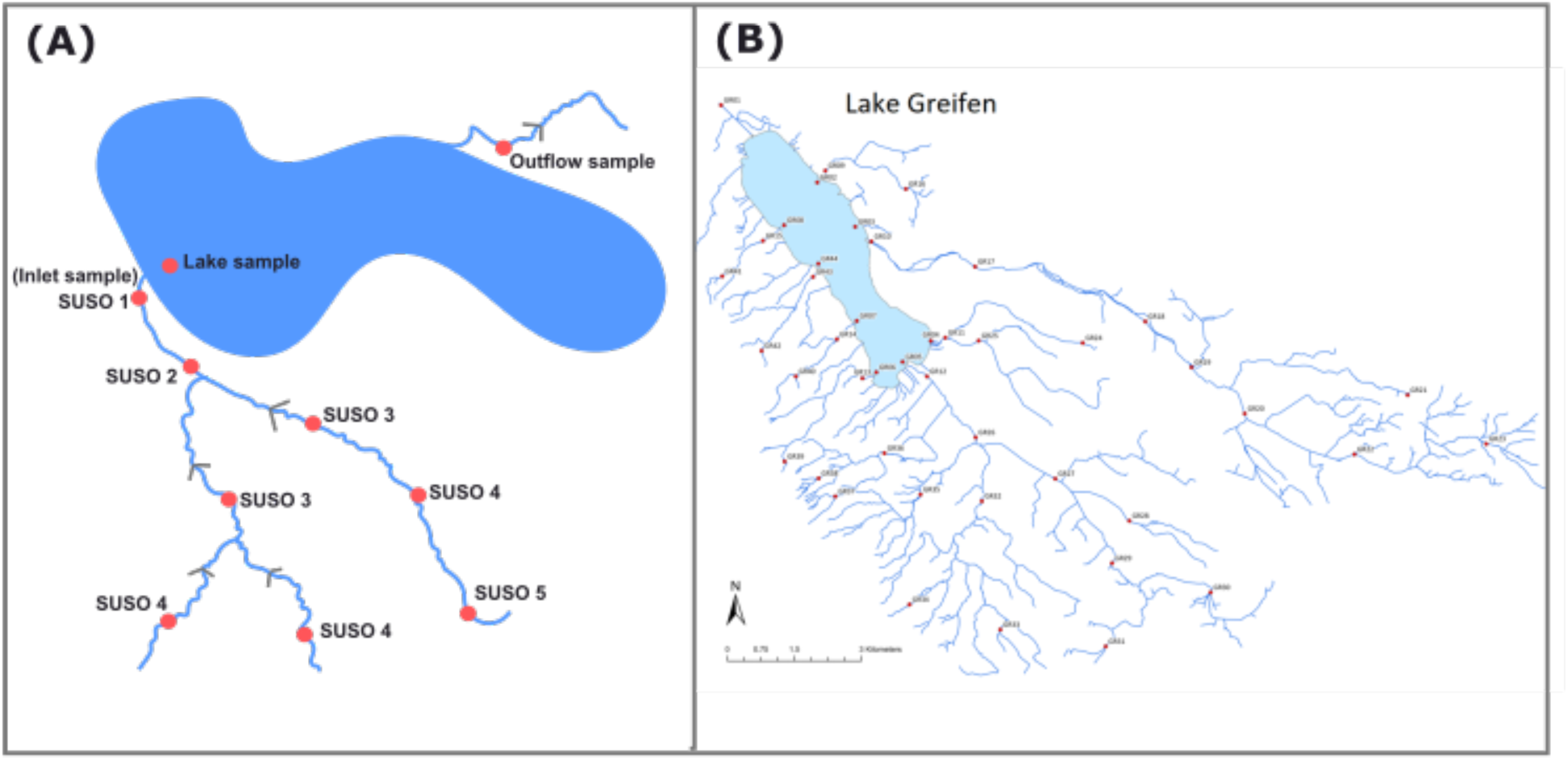
(A) Simplified illustration of sampling design. Red dots indicate sampling locations and gray arrows indicate the direction of water flow. This illustration also provides the Stepwise Upstream Sample Order (SUSO). (B) Map of Lake Greifen watershed with blue lines showing the stream network and red dots indicating sampling locations. Enlarged maps for each of the eight lakes in the study can be found in Supporting Information Figures S1 – S8.

#### SUSO: Stepwise Upstream Sample Order

To calculate eDNA transport distances we annotated the sample sites with Stepwise Upstream Sample Order (SUSO), a stream classification system where numbering starts with “1” from the most downstream point in a stream mainstem and increases at each successive upstream sample point, regardless of confluences. SUSO ensures that every tributary at a confluence receives the same number, while the next upstream point in the same tributary receives a sequentially higher number. The SUSO numbering system is illustrated graphically in Figure 1A. This system can provide a simple, sample-driven hierarchy for studies focused on downstream connectivity and flow pathways.

#### eDNA sampling and state sorting

To sort the eDNA samples into their three states, we developed a new sampling protocol that feeds into the eDNA extraction methodology developed previously (Kirtane et al., 2023; Kirtane & Deiner, 2024). From each site, we collected 100 mL of water and passed the water sample through a 0.22 μm Isopore polycarbonate filter (GTTP02500; Millipore) in 25 mm Swinnex filter holder using 50 mL syringes (Figure 2). The filter holders and syringes were sterilized using a 10% (v/v) dilution of commercially available Javel bleach (≈ 2.5% sodium hypochlorite, ∼2.6% active chlorine) and rinsed with milliQ water, dried, and UV treated for 30 minutes prior to commencing the fieldwork. The membrane-bound and adsorbed states were expected to be accumulated on the filter membrane, while the dissolved state of eDNA was expected to be in the filtrate (Kirtane et al., 2023; Kirtane & Deiner, 2024). The filtrate was collected in 200 mL plastic bottles containing 1 mL of 100x TE buffer. After addition of the filtrate to the concentrated TE, the buffer is diluted to a 1x concentration, reducing enzymatic activity, thus preserving the dissolved eDNA during sample collection and transport. For sites where filters clogged before reaching 100 mL, the volume filtered was noted. After filtering the final volume of the water sample, some air was pushed through the filter to ensure no residual water was remaining in the filter holder. The filter holders were then opened at the field site and carefully transferred into 2 mL lo-bind centrifuge tubes with 750 µL of desorption buffer (0.12 M Na2HPO4, 0.12 M NaH2PO4, 0.1 M EDTA, pH 9). The tubes were brought back to the laboratory or the field station (in the case of Sils and Silvaplana) and shaken for 20 minutes at 21°C at 200 RPM using a shaking incubator allowing all samples ample agitation to desorb the adsorbed eDNA while preventing excessive cell lysis. The tubes were then centrifuged at 13,000 xg for 3 min. At this stage, the adsorbed DNA was expected to be in the supernatant and the membrane-bound DNA in the pellet. The supernatant was then transferred into a new 2 mL lo-bind tube without disturbing the pellet. 750 mL of Longmire lysis buffer (100 mM Tris-HCl (pH 8.0), 100 mM EDTA, 10 mM NaCl, 0.5 % SDS, and 0.8 mg/mL proteinase K) was added to the tube containing the pellet. Then all tubes were frozen at ࢤ20 °C until extraction. The bottles with the filtrate were placed on ice and transported to the laboratory the same day where they were frozen at ࢤ20 °C until further processing. The exception to this were the samples collected from the Sils and Silvaplana watersheds, where the bottles with the filtrate were stored in ice coolers for 7 days and then transported to the ࢤ20 °C freezer in the laboratory.

**Figure 2:**
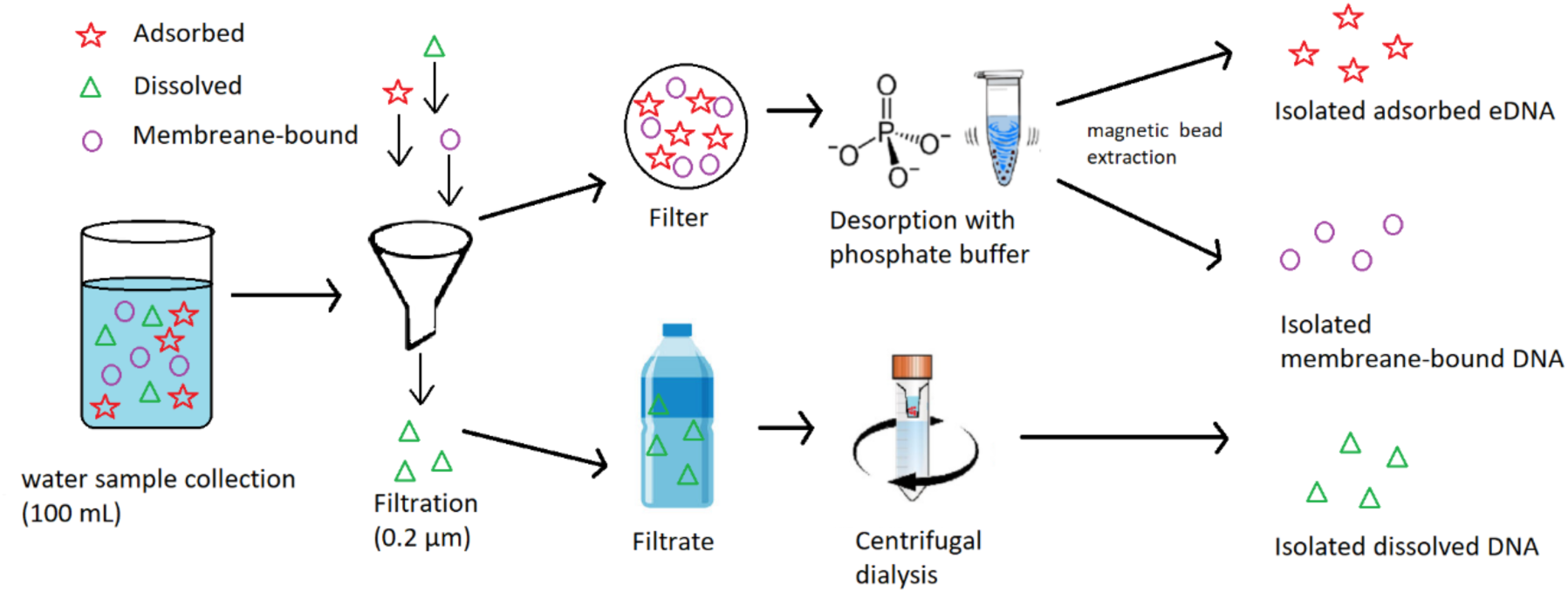
Illustration of sample processing and eDNA state sorting procedure used in this study combining outcomes from (Kirtane et al., 2023; Kirtane & Deiner, 2024). The red stars, green triangles, and purple circles represent adsorbed, dissolved, and membrane-bound states of eDNA, respectively.

Before commencing most sampling expeditions, a full process negative control was performed, where 100 ml of Milli-Q water was processed identically to the field samples in the field. A total of five full process negative controls were included in the study, which were state sorted into three states resulting in fifteen full process negative controls sequenced. Additionally, fifteen extraction controls (five per state) and twelve PCR negative controls were used to monitor contamination at different steps of the workflow for a total of 42 negative controls.

We collected various co-variables alongside the water samples. These included the coordinates of the samples, the type of substrate at the sampling location, the date, time, and water matrix characteristics including pH, temperature, total dissolved solids, and electrical conductivity using a multiprobe (HI9813-6, Hanna Instruments, Woonsocket, RI, USA) (Table S1, Figure S9).

### DNA extraction

#### Membrane-bound eDNA state

The tube containing the membrane-bound eDNA, i.e., the tube with the pellet in 750 µL of Longmire lysis buffer with proteinase K was thawed and incubated overnight in a shaking incubator at 55 °C. The lysate was then transferred into 2 mL 96-deep well plates and extracted using a magnetic bead extraction protocol programmed on an Opentrons liquid handling robot. The protocol entailed first placing the 96-deep well plate with the lysate on the Opentrons magnetic module. Then, 600 µL of binding matrix containing 500 µL of Environmental Hybridization buffer (For 500mL: 1 g DTT, 72.5 g NaCl, 125 g PEG 8000, 500 μL 0.5M EDTA) and 100 µL 20 % Sera-Mag SpeedBead Carboxylate modified magnetic beads (GE Healthcare, IL, USA) in TE buffer was added into each of the wells. The mixture was then pipette mixed ten times and incubated at room temperature for 10 minutes. The magnetic module was engaged, allowing the magnetic beads to migrate to the magnet for 30 minutes. During this time, the mixture was gently mixed by pipetting again ten times to ensure all beads could encounter the magnetic field and migrate towards the magnet. Once beads migrated to the magnet, indicated by the clarity of the solution, the supernatant was gently pipetted out and discarded into the waste container. Next, the magnetic module was disengaged, and 1.5 mL of 80% ethanol was added to each well and pipetted gently to wash off any remaining buffer. The magnetic module was re-engaged, and the beads were once again allowed to migrate to the magnets. The supernatant ethanol was then discarded. This process was repeated one more time. After the second ethanol wash, the magnetic beads were allowed to air dry for 1.5-2 hours to ensure evaporation of residual ethanol. The magnets were disengaged and 200 µL of TE buffer was added to elute the DNA from the magnetic beads and mixed by pipetting five times. The magnetic module was re-engaged for the last time, allowing the beads to migrate to the magnets. The supernatant was then transferred to a 96-well PCR plate, which was immediately sealed and frozen at ࢤ20 °C until further processing.

#### Adsorbed eDNA state

The adsorbed DNA was extracted using the same magnetic bead protocol using an Opentrons liquid handling robot. However, instead of 750 μL of pellet in the lysis buffer in the first step, 750 μL of the desorption buffer containing the adsorbed state of eDNA was added.

#### Dissolved eDNA state

Dissolved state eDNA samples were extracted by centrifugal dialysis using Amicon® tubes (Kirtane & Deiner, 2024). Samples were first thawed in an incubator at 55 °C. Approximately 15 mL of the sample was then added to a 10,000 MWCO Amicon® ultrafiltration tube and centrifuged (Eppendorf Centrifuge 5804 R) at 4200 xg at 23 °C for 15 minutes. The flow-through was discarded, and the tube was refilled with another 15 mL of sample. This was repeated until the full 100 mL had been concentrated. During field sampling, a white precipitate formation was noticed in 52 out of 228 dissolved state samples. If there was no precipitation, 1 mL of 1x TE buffer was added to wash the dialysis membrane and centrifuged again to obtain approximately 250 µL of DNA, which was stored at ࢤ20 °C in 1.5 mL LoBind® Eppendorf tubes. The precipitated samples were treated similarly but did not concentrate down to 250 μL, but instead around 1-1.5 mL. This 1 – 1.5 mL of concentrate was then transferred to 2 mL LoBind® Eppendorf tubes, which were then centrifuged to pellet the precipitate and the supernatant transferred to new tubes with the rest of the samples. After purification and concentration, all dissolved state concentrated samples underwent ZYMO inhibitor removal (OneStep PCR Inhibitor Removal Kit) according to the manufacturer’s protocol and were then stored at ࢤ20 °C in 96-well plates.

### DNA quantification and metabarcoding analysis

After DNA extraction, all eDNA samples were quantified using the Spark® 10M Multimode Microplate Reader (Tecan, Männedorf, Switzerland) with Qubit HS Kit reagents. The quantification was conducted in 384-well plates with a reaction volume of 50 µL per well, consisting of 2 µL DNA and 48 µL reaction mix (199:1 ratio of iQuantTM buffer to HS iQuantTM reagent dye ratio (Qiagen, Hilden, Germany)). All samples were analyzed in duplicate. Each quantification plate included a six-point standard curve (10 ng/µL, 5 ng/µL, 1 ng/µL, 0.5 ng/µL, 0.1 ng/µL, 0 ng/µL) in duplicate to create a linear regression to calculate the eDNA concentration of the samples.

#### PCR and library preparation

Metabarcoding analysis for the COI marker for metazoans was performed using the Sauron-S878 forward primer (Rennstam Rubbmark et al., 2018) with reverse primer HCO2198 (Folmer et al., 1994). The metabarcoding for the ITS marker for plants was performed using ITS2 primers (Chen et al., 2010). All primer sequences can be found in Table S2. The same PCR library preparation protocol was performed for both COI and ITS markets. The first involved a boost PCR followed by a second PCR using the single-tube protocol described in Rennstam Rubbmark et al. (2018) to attach multiplex identifiers (MIDs) and Illumina sequencing adapters within one reaction (Table S2).

Boost PCR was conducted in 384-well plates with 4 replicates per sample where each well contained 5 µL master mix (2x Multiplex PCR Kit (QIAGEN)), 0.25 µL RSA ([20 mg/ml] New England Biolabs), 0.4 µL boost primer mix [10 µM], 1.5 µL DNA and 2.85 µL molecular grade water to get a reaction volume of 10 µL. DNA was transferred from 96-well plates to the reaction mixture in the 384-well PCR plates Opentrons OT-2 Liquid Handler. The PCR was conducted on a Labcycler (SensoQuest GmbH, Göttingen, Germany) with an initial incubation step at 95 °C for 15 minutes, followed by 25 cycles with denaturation at 95 °C for 30 seconds, annealing at 56 °C for 90 seconds, and extension at 72 °C for 60 seconds and a final incubation at 72 °C for 5 minutes. After boost PCR, replicates from the 384-well plate were pooled into a 96-well plate using the Liquid Handling Station (BRAND Scientific, Essex, CT, USA). After PCR, clean-ups were performed using 1:0.8 PCR product to Ampure XP magnetic beads (Beckman Coulter, Brea, CA) to select the target amplicon.

The cleaned boost PCR product was then further amplified using the single tube protocol in 96-well plates with one replicate per sample (Rennstam Rubbmark et al., 2018). Each reaction contained 10 µL master mix (2x Multiplex PCR Kit (QIAGEN)), 0.5 µL RSA ([20 mg/ml] New England Biolabs), 0.3 µL universal primer mix [10 µM], 3 µL index primer mix [10 µM], 3 µL of the PCR product, and 3.2 µL molecular grade water to get a final reaction volume of 20 µL. Individual index primers were added using the OT-2 Liquid Handler. Single-tube PCR cycling had an initial incubation step at 95 °C for 15 minutes, followed by 10 cycles with denaturation at 95 °C for 30 seconds, annealing at 56 °C for 90 seconds, and extension at 72 °C for 60 seconds. This was followed by 10 cycles with denaturation at 95 °C for 30 seconds, annealing at 45 °C for 90 seconds, and extension at 72 °C for 60 seconds, followed by a final incubation at 72 °C for 5 minutes. This was followed by another magnetic bead cleanup as described above. After cleanup, a random subset of samples were visualized on 1.5% agarose gels to ensure amplification of the desired amplicon size and the success of the magnetic bead cleanup. ITS marker bands of the expected size were absent in most adsorbed and dissolved samples; therefore, only membrane-bound samples were used for sequencing the ITS marker.

After clean-up, the PCR products were quantified using Spark® 10M Multimode Microplate Reader with Qubit BR kit reagents as described above in the DNA quantification section. Based on the result of the plate reader measurement, the PCR products were pooled at equimolar concentrations using a pipetting robot (Liquid Handling Station, BRAND Scientific, Essex, CT, USA) to ensure an even sequencing depth across samples. The PCR products were pooled into one of two pools, one for each lane of the sequencing flow cell. After pooling, a second clean-up was performed with Ampure XP beads (Beckman Coulter, Brea, CA) at a ratio of 1:0.7 PCR product to Ampure XP beads for all pools.

To remove heteroduplexes, a reconditioning PCR was conducted on the pooled library using eight replicate reactions with 12.5 µL Kapa HiFi HotStart ReadyMix (2x Kapa Biosystems, Roche), 1 µL of P5 primer [10 µM], 1 µL of P7 primer [10 µM], 3 µL of cleaned index PCR product, and 6.5 µL of water to get a reaction volume of 25 µL. Reconditioning PCR was conducted with an initial incubation step at 95 °C for 3 minutes, followed by 7 cycles with denaturation at 98 °C for 20 seconds, annealing at 62 °C for 15 seconds, and extension at 72 °C for 30 seconds. This was followed by a final incubation at 72 °C for 1 minute. Subsequently, another clean-up with Ampure XP beads (Beckman Coulter, Brea, CA) at a ratio of 0.7:1 reconditioning PCR product to Ampure XP beads was conducted with each pool. The pooled libraries were then sequenced by the Functional Genomics Center Zurich (FGCZ) using the Illumina NovaSeq 6000 SP 2x250bp flow cell Reagent Kit v1.5 with 20% PhiX.

### Bioinformatics

The bioinformatics processing of sequencing data was performed separately for the COI and ITS amplicons using different pipelines optimized for each marker.

#### COI Amplicon Processing

For the bioinformatics of COI amplicon sequences, UPARSE (OTU) and UNOISE (ASV) workflows were used to process the Illumina paired-end sequence reads (Edgar, 2010, 2013). Samples with low read counts (<1000) were initially removed, leaving 713 samples (92.8%). Then the COI-related reads were extracted. Given that we used custom sequencing primers that match the target primer site, our reads had no primer sequences as part of the amplicon. Thus, we could not find loci-specific reads and determine the orientation of the amplicon by searching the primer sequence. Instead, we used a sequence alignment approach (i.e., usearch_global) to find COI-related reads. We created a reference for sequence alignment searches using the MIDORI2 CO1 (GB259) database (Leray et al., 2022). The first step was to perform an in-silico PCR to obtain potential amplicons. The next step was to cluster the sequence using a 10% similarity radius. The 12,520 unique COI sequences were used to capture COI-related reads for the dataset. After extraction, we again filtered out samples with fewer than 1000 reads. We merged the read pairs and quality-filtered them. The UNOISE3 and UPARSE2 algorithms were used to cluster the reads. Singletons (abundance threshold = 2) were removed. UNOISE3 applies error correction and zero-radius clustering to find ASVs that can be considered ASVs. A minimum abundance threshold of 7 was used. We used USEARCH: SINTAX together with the MIDORI2 COI reference (GB242, Leray et al., 2022) and applied a confidence threshold of 0.75 to predict taxonomic associations.

#### ITS Amplicon Processing

The bioinformatic analysis was conducted using the DADA2 pipeline in R Studio (Callahan et al., 2016; R Core Team, 2021) to process ITS paired-end sequence reads. The analysis followed the ITS-specific workflow from the DADA2 tutorial (v1.29.0). Due to the long amplicon length of the ITS marker, the paired-end reads could not be merged, and only forward reads were retained for further analysis. Reads containing ambiguous bases (Ns) were removed, and a quality filtering step was applied to exclude low-quality reads (maximum expected errors = 2, truncation length = 251 bp). Following quality filtering, error rates were learned using the ‘learnErrors’ function, and denoising was performed using the dada function to infer ASVs. Chimeric sequences were identified and removed using the ‘removeBimeraDenovo’ function, applying a consensus-based approach. After quality control, an ASV table was generated. To further refine clustering, ASVs were processed using the Swarm v3 algorithm (Mahé et al., 2015), which performs single-linkage clustering to generate biologically meaningful sequence clusters. Taxonomic classification was performed using the VSEARCH: SINTAX (Rognes et al., 2016) with the PLANiTS ITS2 reference database (Banchi et al., 2020). A confidence threshold of 0.75 was applied to assign taxonomic identities.

### Data analysis

All data analysis was conducted in R Studio using the ‘phyloseq’ package (McMurdie & Holmes, 2013; R Core Team, 2021). ASVs detected in any of the negative controls were removed from the entire dataset prior to downstream analyses. All ASVs found only in one sample in the entire dataset were removed. All samples were rarified at 10,000 reads and samples below 10,000 reads were removed from the dataset (Figure S2). The taxonomic identity of the ASVs was updated by synchronizing it with the GBIF (Global Biodiversity Information Facility) backbone using the R package ‘rgbif’ (Chamberlain et al., 2017). While primary analyses were conducted at the ASV level, species-level assignments were referenced solely to illustrate selected examples in the discussion, without influencing the statistical results. Analyzing transport with ASVs is a conservative approach that ensures that the same sequence was tracked along the catchment, not the inference of species that could be comprised of several ASVs (Tsuji et al., 2020).

To evaluate variations in the eDNA information captured in the three states, a subset of samples was created that only included those where all three states were successfully sequenced (n = 138). These samples were compared based on their DNA yield, ASV richness, Shannon-Wiener index value, and the number of unique ASVs within each state. Statistical analysis involved the Friedman test, grouped by sampling site, followed by the Wilcoxon signed-rank post hoc test to determine differences in DNA yield, ASV richness, and Shannon-Wiener index. The community composition was visualized at the “phylum” level for COI and “class” level for ITS taxa.

The transport analysis was divided into two components: 1) transport from upstream sites to downstream sites, and 2) transport from streams into lakes. For eDNA transport within streams, flow-connected stream samples (i.e., upstream and downstream samples) were identified. The difference in SUSO values between flow-connected samples was used as a proxy for distance. Percent shared AVSs between flow-connected sample pairs was calculated, defined as the number of shared ASVs divided by the total number of unique ASVs observed across both samples, multiplied by 100. Spearman correlation was used to assess the relationship between percent shared richness and SUSO difference.

To investigate eDNA transport dynamics from inlets to lakes, we compared ASV richness across sampling positions (up-stream, inlet, lake, and outflow) using the Kruskal-Wallis test, followed by Dunn’s post-hoc test with Benjamini-Hochberg correction. We also investigated the impact of the distance between flow-connected inlet and lake sampling sites on the shared richness of taxa between them. Exact distances between sampling sites were calculated using the ‘rivnet’ R package. Linear regression analysis was performed to investigate the significance of the relationship.

Differences in community composition among stream, inlet, lake, and outflow samples were visualized using NMDS plots based on Jaccard distance, with environmental variables (temperature, pH, total suspended solids, and electrical conductivity) overlaid to assess their influence across eDNA states and markers. ASVs present in fewer than three samples were excluded to reduce spurious overlap and improve distance estimation. Outliers in ordination space were removed based on a 1.5× IQR threshold along each NMDS axis to improve visualization and vector fitting. Community differences among stream, inlet, lake, and outflow samples were tested using pairwise PERMANOVA on Jaccard distances with Bonferroni-adjusted p-values. The influence of environmental variables (temperature, pH, total suspended solids, and electrical conductivity) on community composition was evaluated by fitting vectors to the NMDS ordinations using permutation-based multivariate regression (‘envfit’). Pairwise PERMANOVA was performed using the ‘pairwiseAdonis’ package, while ‘phyloseq’ was used to calculate ASV richness, Shannon-Weiner index, and Jaccard distance. All plots were created using ‘ggplot2’ (Wickham et al., 2016). The Friedman test, Kruskal-Wallis’s test, Dunn’s test, and Wilcoxon signed-rank tests were performed using the ‘stats’ and ‘rstatix’ packages. Portions of the R code used for data analysis were debugged and refined with the assistance of ChatGPT (GPT-4o, OpenAI).

## Results

For the COI amplicon sequences, eleven negative controls (four full-process, six extraction, and one PCR control), out of a total of forty-two, had detectable sequences with a total of 39 ASVs, which were removed from the entire dataset. After rarifying to a threshold of 10,000 reads, 18 samples from the dissolved state and 2 samples from the membrane-bound state were removed, resulting in 573 samples in the dataset (Figure S10A). This contained a total of 29,736 ASVs, most of which (74.7%) did not have a taxonomic identity match in the reference database. Taxonomic assignments (confidence threshold = 0.75) resulted in 25.3 % of sequences assigned to a phylum, 18.4 % to a class, 15.3 % to an order, 14.1 % to a family, 13.4 % to a genus, and 10.0 % to a species.

For the ITS amplicon sequences, all fourteen negative controls (five full-process, six extraction, and three PCR controls) had 384 ASVs which were removed from the entire dataset. After rarifying to 10,000 reads, none of the samples had to be removed resulting in a total of 221 samples (Figure S10B) containing a total of 7246 ASVs. Taxonomic assignments (confidence threshold = 0.75) resulted in 28.8 % of sequences assigned to a phylum, 27.5 % to a class, 27.4 % to an order, 27.2 % to a family, 26.6 % to a genus, and 9.19 % to a species.

### eDNA State-Dependent DNA Yield and Diversity

The number of samples successfully sequenced over the sequencing depth threshold depended on the eDNA state. For the COI marker, the samples from the dissolved state had the fewest samples (n = 144) successfully sequenced, followed by adsorbed (n = 214) and membrane-bound (n = 215) states, showing a state-dependent success rate for amplification and sequencing. To explore variations between the states, only sites that had successful amplification represented by all three states (n = 138) were used for visualizations and statistical analyses in this section. No samples were removed for the eDNA transport analyses as these analyses compare samples within the same state in the following sections. For the ITS marker, only the membrane-bound DNA state could successfully be amplified and sequenced.

#### DNA yield

DNA yield from the three extracted states shows effective extraction of all three DNA states (Figure 3A). However, DNA yield was significantly different across adsorbed, dissolved, and membrane-bound states (Figure 3A, Friedman chi-squared = 220.60, df = 2, p-value < 2.2e-16). Pairwise Wilcoxon signed rank test further identified significant differences between the DNA yield of each state showing that the dissolved state had significantly higher DNA yield compared to membrane-bound (p.adj = 4.9e-14) and adsorbed states (p.adj < 2e-16), while the adsorbed state also had the DNA yield significantly lower than the membrane-bound state (p.adj < 2e-16) (Figure 3A).

**Figure 3:**
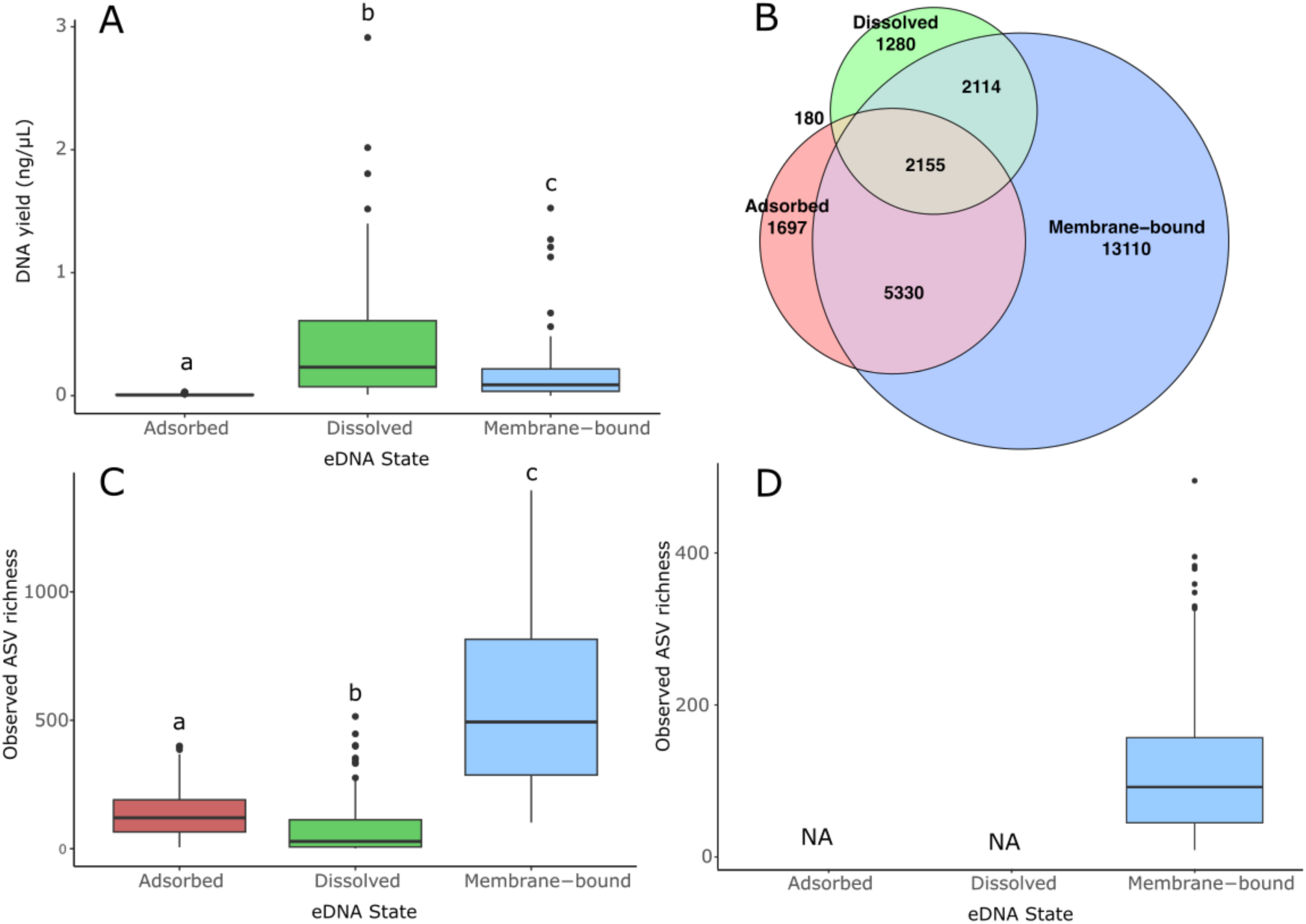
(A) Boxplot showing the DNA yield; (B) Venn diagram showing the number of shared and unique metazoan (COI marker) ASVs; (C) observed richness of metazoan (COI marker) ASVs. This analysis only includes samples that were successfully sequenced in all three states using the COI marker (n = 138). (D) observed richness of plant (ITS marker) ASVs. The letters above the bar plots (a,b,c) indicate statistically significant differences.

#### Alpha richness and diversity

For the COI marker, the mean ASV richness captured was significantly different between each eDNA state (Figure 3C, Friedman chi-squared = 294.86, df = 2, p-value < 2.2e-16). Pairwise Wilcoxon signed rank test showed that the membrane-bound state has the highest mean ASV richness (Figure 3C), significantly higher than adsorbed (p.adj < 1.42e-23) and dissolved (p.adj 1.45e-23) states, while the dissolved state also had significantly lower ASV richness than the adsorbed state (p.adj = 6.27e-6) (Figure 3C). The membrane-bound state also had a significantly higher Shannon-Wiener index value indicating high mean richness and evenness within this state compared with the other two states (Figure 3D; Friedman chi-squared = 108.81, df = 2, p-value < 2.2e-16). The dissolved state had the lowest Shannon-Wiener index value compared with adsorbed (Figure 3D; p-adj < 5.42e-11) and membrane-bound eDNA states (Figure 3D; p-adj < 7.44e-19). The Venn diagrams show that the majority of the ASVs captured were unique to one of the three states (62.6 %) with 6.5 % unique to the adsorbed state, 4.6 % unique to the dissolved state, and 50.8 % unique to the membrane-bound state (Figure 3B). Only 8.4 % of all ASVs were shared among all three states despite originating from the same set of water samples (Figure 3B).

Each of the three states had a variable COI community composition despite originating from the same samples. However, the community composition was more similar in the adsorbed and membrane-bound state than in the dissolved state (Figure 4). For instance, the dissolved state had a much greater proportion of its diversity, almost 50 % of all taxa in stream samples, from the phylum Ascomycota in stream samples and Haptophyta in lake samples, while phyla such as Amoebozoa, Oomycota, Ochrophyta, and Rotifera had a higher proportion of taxa captured in adsorbed and membrane-bound states compared to the dissolved state (Figure 4).

**Figure 4:**
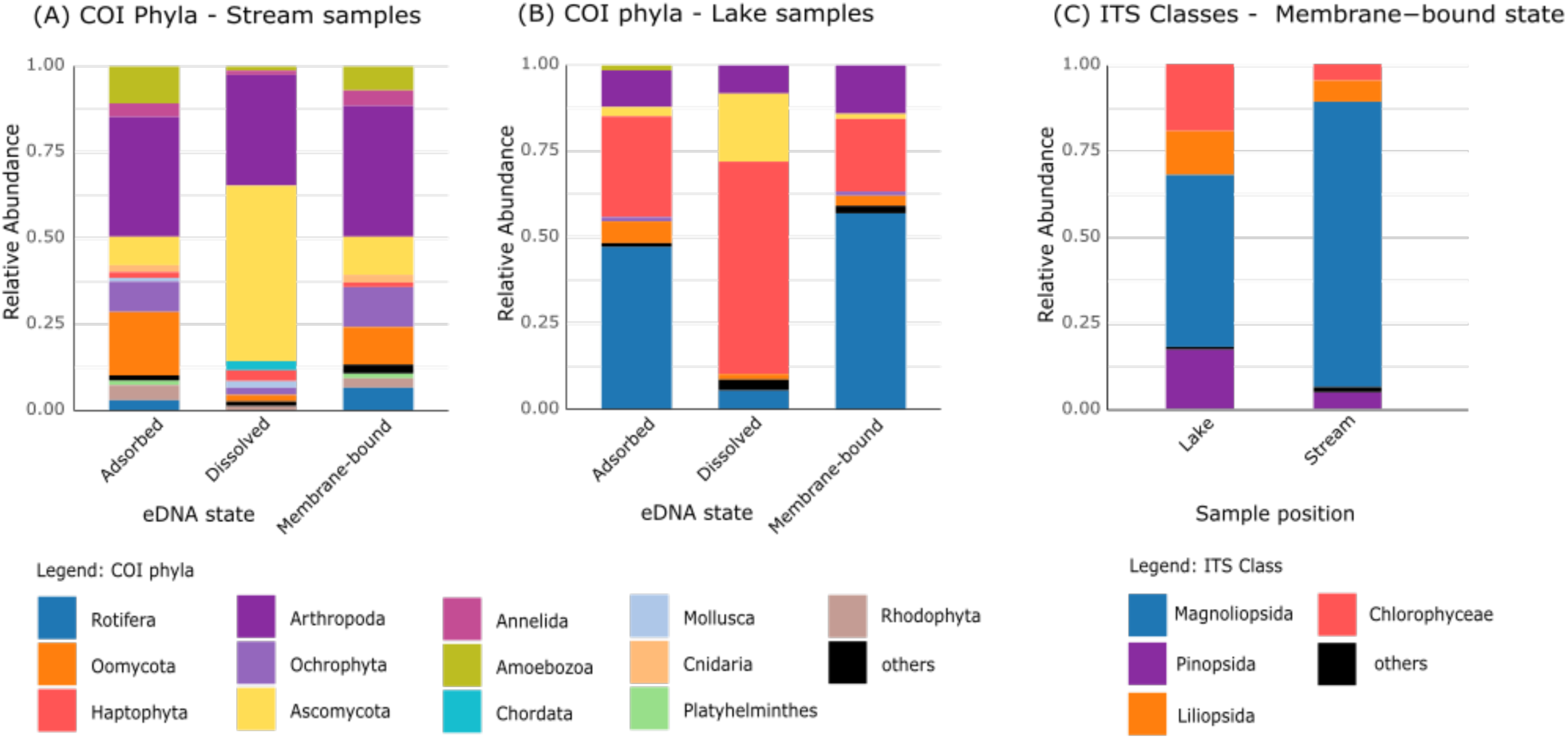
Stacked bar plots (A and B) show the relative abundance of phyla for the COI marker in different eDNA states in (A) stream samples, and (B) lake samples. (C) Classes for the ITS marker for the membrane-bound state in stream and lake samples. Taxa with relative abundance below 1% are grouped as “others.”

The COI community composition was different between the stream and the lake samples. While the same phyla were shared between the stream and the lake samples there were some key differences. Groups such as Haptophyta and Rotifera represented a greater proportion of the captured taxa in the lake samples compared to the stream samples, while Ascomycota, Arthropoda, and Ochrophyta had a greater proportion in the stream samples across all states (Figure 4).

### Downstream Transport of eDNA States from Headwaters to Lake Inlets

The transport behavior of eDNA through streams was impacted by the state of eDNA. Flow-connected samples (i.e., two samples along the same stream) were identified and resulted in a total of 207 pairs for the membrane-bound state, 203 pairs for the adsorbed state, and 99 pairs for the dissolved state with the COI marker, and 217 pairs for the membrane-bound state using the ITS marker.

The percentage of shared ASV richness of flow-connected sample pairs plotted along the difference of SUSO between them showed a general pattern of decreasing shared richness with an increase in SUSO difference (Figure 5). This trend was expected as SUSO is a proxy for distance, and with increased distance, the eDNA community captured can change as the eDNA degrades over time and new eDNA may get added to the stream. The Spearman correlation analysis reveals negative associations between SUSO difference and percent shared ASV richness across the membrane-bound state, indicating that as samples become more spatially separated (higher SUSO difference), they tend to share fewer taxa. This relation was statistically significant for both the COI marker (*Spearman’s rho* = ࢤ0.27, *p* = 6.49e-05) and the ITS marker (*Spearman’s rho* = ࢤ0.22, *p* = 1.12e-03). This relationship did not have a statistically significant correlation for the adsorbed (*Spearman’s rho* = ࢤ0.09, *p* = 0.19) and dissolved states (*Spearman’s rho* = ࢤ0.08, *p* = 0.45). For the COI marker, this trend was particularly strong and statistically significant in the membrane-bound state in specific sites, including Aegari watershed (*Spearman’s rho* = ࢤ0.60, *p* = 0.003), Baldegg watershed (*Spearman’s rho* = ࢤ0.62, *p* < 0.001), and Greifen watershed (*Spearman’s rho* = ࢤ0.37, *p* = 0.005), suggesting a consistent spatial decay of richness similarity in membrane-bound DNA state across watersheds (Table S3, Figure S12, Figure S14). Watershed-level data figures and Spearman’s correlation table faceted by watershed, state, and marker can be found in Figure S12 and Table S3, respectively.

**Figure 5:**
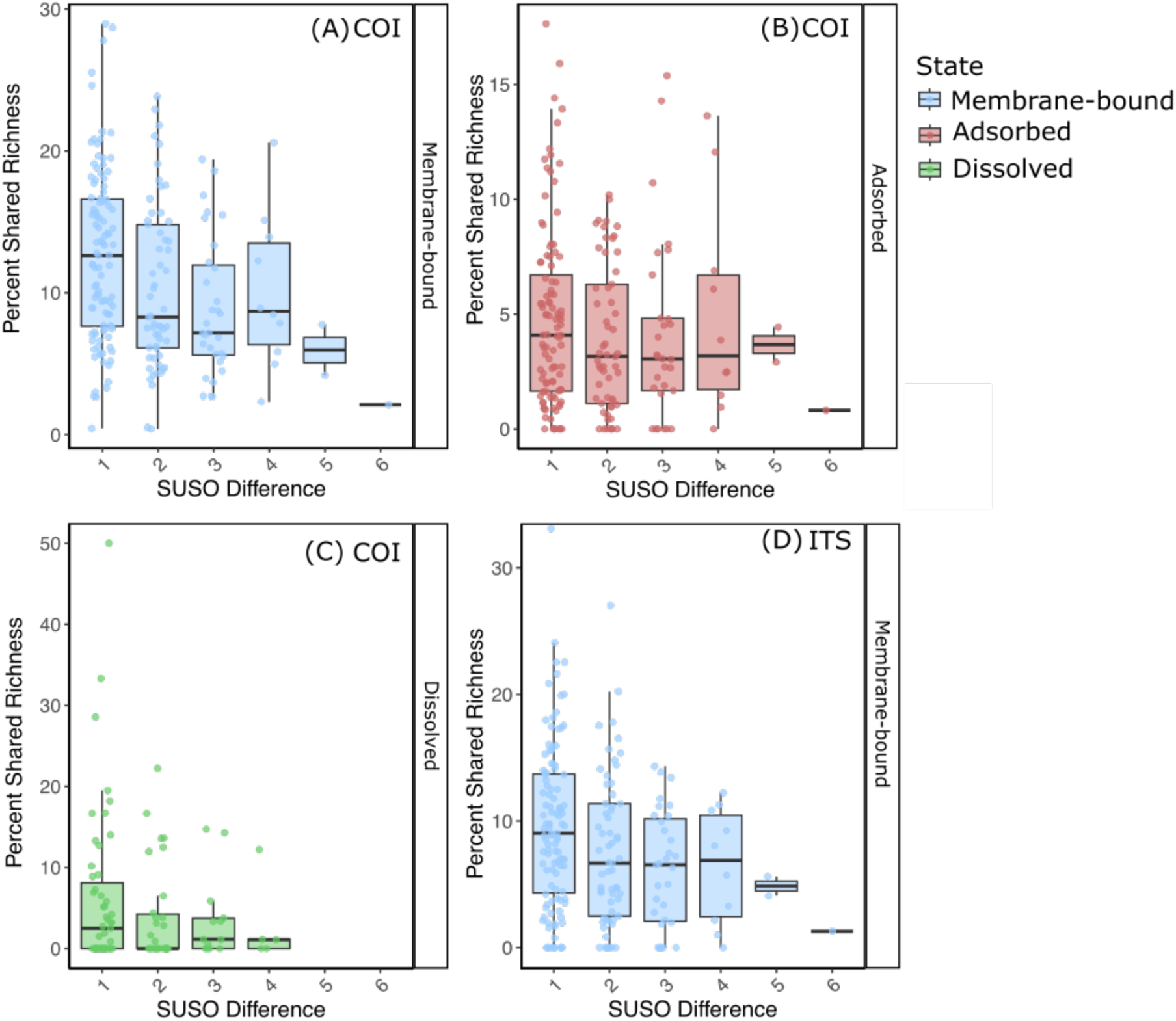
Percent shared richness (observed ASV richness) between flow-connected sample pairs over SUSO difference for the COI marker (A, B, C representing membrane-bound, adsorbed, and dissolved states, respectively) and the ITS marker (D representing the membrane-bound state). The absence of SUSO differences 5 and 6 in the dissolved state is due to the unsuccessful sequencing of samples.

### Comparative Biodiversity in Stream Inlets and Lakes

To examine eDNA transport from streams to lakes, we categorized samples into four groups: stream, inlet, lake, and outflow. Stream samples were collected from headwater streams, excluding inlets, which were defined as the final sampling point before a stream entered a lake. Stream sampling points were spaced 1–3 km apart, whereas inlet samples were located 10–750 m from the lake. Lake samples were taken directly from the surface of the lake 2-50 m from the mouth of the stream.

In the COI dataset, ASV richness did not differ significantly among sampling locations for the adsorbed DNA (Figure 6A). For the dissolved state, lake samples showed higher ASV richness than stream and inlet samples (Figure 6B). In the membrane-bound state, stream and inlet samples had significantly higher richness than lake and outflow samples (Figure 6C). In the ITS dataset, stream samples exhibited significantly higher ASV richness than lake samples (Figure 6D). Full statistical results from the Kruskal–Wallis and Dunn’s post hoc tests are provided in Tables S6 and S7.

**Figure 6.**
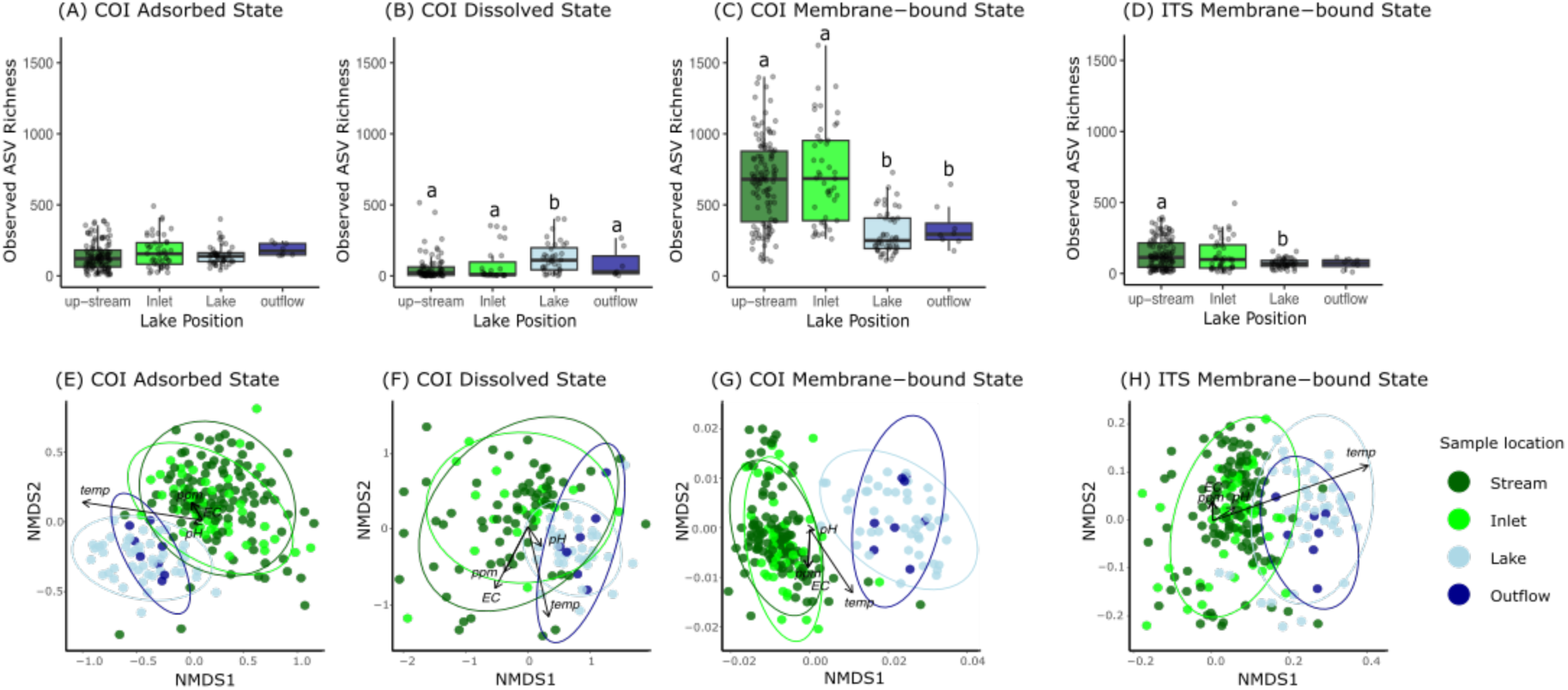
ASV richness based on the COI marker for (A) adsorbed, (B) dissolved, and (C) membrane-bound DNA, and based on the ITS marker for (D) membrane-bound DNA. Samples were collected from streams, inlets, lakes, and outflows. Different letters above the box plots (a, b) indicate statistically significant differences between the means of groups. Panels (E–H) show NMDS plots with Jaccard distances of ASV for different eDNA states and markers. Each panel shows community composition for (E) adsorbed, (F) dissolved, and (G) membrane-bound states using the COI marker, and (H) membrane-bound state using the ITS marker. Arrows represent environmental variables (temperature (temp), pH, electrical conductivity (EC), and total suspended solids (ppm) significantly correlated with community structure (p ≤ 0.05). Arrow lengths were scaled independently for each plot for visual clarity and are not comparable across plots. Statistical parameters for these relationships can be found in Table S8.

NMDS plots showed that stream and inlet samples clustered together, while lake and outflow samples formed a separate cluster – a consistent pattern across all states and both COI and ITS markers (Figure 6E-H). PERMANOVA results confirmed no significant difference in community composition between stream and inlet samples, nor between lake and outflow samples (Table S4). Notably, the explanatory power of these tests (R² values) increased when comparisons were restricted to sampling locations within the same lake watershed, likely due to reduced environmental heterogeneity and pooling effects (Figure S11, Table S5). A weak negative relationship was observed between the distance separating inlet and lake sample pairs and their shared richness, but only in the membrane-bound state (R2 = 0.13, p = 0.02; Figure S15).

### Environmental vector fitting analysis revealed state-specific drivers of community structure

Environmental vectors overlaid on NMDS ordinations revealed that temperature and pH were strongly aligned with gradients in community composition across multiple eDNA states, while total suspended solids and electrical conductivity showed weaker alignment. Nevertheless, all fitted environmental variables were statistically significant (p = 0.001; Figure 6, Table S8). In all ordinations, the temperature vector consistently pointed from stream and inlet samples toward lake and outflow samples, suggesting that community composition shifts along a thermal gradient (Figure 6). In membrane-bound state samples, temperature (R² = 0.5), electrical conductivity (R² = 0.26), and total suspended solids (R² = 0.26) showed strong and significant associations with the NMDS ordination (p = 0.001), while there was a weaker association with pH (R² = 0.06) for the COI marker (Table S8). In contrast, both dissolved and adsorbed samples showed a strong correlation with temperature (R² = 0.28 and R² = 0.17, respectively) and weaker (R² < 0.1) but still significant (p < 0.01) associations for pH, electrical conductivity, and total suspended solids (Table S8). In the ITS membrane-bound state, environmental variables showed weaker overall correlations, with the strongest association observed for temperature (R² = 0.15), and all vectors remaining significant at p < 0.01 (Table S8).

## Discussion

When the concept of eDNA states was first proposed, it was hypothesized that different states might exhibit distinct behaviors in aquatic environments, potentially influencing both the biodiversity information they carry and their patterns of transport (Jo & Yamanaka, 2022; Mauvisseau et al., 2022; Nagler et al., 2022). This study verifies these hypotheses, showing that the biodiversity information collected depends on the eDNA state, and the eDNA state influences its transport behavior. The results indicate that the three eDNA states – adsorbed, dissolved, and membrane-bound – carry different biodiversity information, with the membrane-bound eDNA state containing the majority of unique ASVs for COI (metazoan) amplicons. Furthermore, membrane-bound state was the only detectable state for ITS (plant) amplicons. However, the other states also contained unique COI sequences not found in the membrane-bound state. Our findings suggest that past eDNA studies, which primarily capture membrane-bound eDNA, have likely reflected most of the taxonomic diversity present in water samples. However, capturing and analyzing all three states from water samples provides a more complete picture of the biodiversity information within an eDNA sample.

The state of eDNA also impacted its transport behavior. Generally, the percentage of shared ASVs in flow-connected samples decreased with increased distance between them. This relationship was statistically significant only for the membrane-bound state for both COI and ITS markers, but was not statistically significant for the adsorbed or the dissolved states. Thus, membrane-bound DNA is being transported, whereas the other two states show little evidence of being transported. While eDNA in the membrane-bound state was transported to the inlet, lake surface samples collected 2-10 meters from the inlet showed a marked turnover in ASV composition. Of the measured water matrix characteristics, temperature seems to be the main driver for this pattern. Further sampling scenarios need to be tested to determine the fate of transported eDNA in lakes and determine the optimal sampling strategies to capture it.

### Richness and Diversity Differences Among eDNA States

This large-scale field study with state-specific metabarcoding analysis showed that the biodiversity information collected is dependent on the eDNA state captured. While the adsorbed and dissolved states of eDNA were comprised of sequences unique to their respective states, the membrane-bound DNA had the majority (81.7 %) of unique sequences (Figure 3B). Most eDNA diversity may reside in the membrane-bound state due to biological factors—such as how DNA is shed or persists within cells—though the exact processes are not yet well understood. Another explanation could be the biases caused by the eDNA state sorting workflow which extracts the membrane-bound state with higher efficiency than other states (Kirtane et al., 2023).

Most eDNA studies use sampling and extraction workflows such as filtration followed by Qiagen Blood and tissue kit extraction (Takahashi et al., 2023), enriching for the membrane-bound eDNA state. Most eDNA from eukaryotic organisms may be released and persist in the membrane-bound state in the water column sourced through molt, feces, spawn, etc. Size fractionation studies revealed that dissolved eDNA accounted for less than 10 % of the total eDNA from mackerel and sardine (Sassoubre et al., 2016). Similarly, the percentage of dissolved eDNA from common carp in the filtrate i.e., dissolved state, was significantly lower compared to its eDNA collected on filters, i.e., membrane-bound state (Turner et al., 2014). A metabarcoding study in marine waters showed that most of the amplifiable eDNA from eukaryotic species was detected in filters with a pore size of 5–10 µm, which captured membrane-bound DNA, while the dissolved state in the filtrate had the lowest species richness among all the size fractions investigated (Power et al., 2023). These findings corroborate the results of this study that the majority of eDNA diversity is harbored and persists within the membrane-bound state. Given that most published studies for the entire field of aquatic eDNA do focus on filtration, it is confirmed here that filtration is the most robust sampling method to detect the majority of eDNA in aquatic systems.

The dissolved state of eDNA is hypothesized to be removed by rapid degradation compared to membrane-bound eDNA via extracellular enzymes and photochemical oxidation leading to a lower sequence richness detected (Lindahl, 1993; Torti et al., 2015). However, the fluorometry measurements showed that the dissolved state had the highest eDNA yield of all three states (Figure 3A). One might initially expect this to result in a greater number of detectable taxa, but our findings show that the dissolved state of eDNA had the minority (22.1 %) of total sequences (Figure 3B) supporting the idea that dissolved eDNA is rapidly degraded and becomes increasingly difficult to sequence. Similar to the results of this study, a size fractionation study also reported dissolved eDNA having the highest eDNA concentration as measured by fluorometry, but did not have equivalent PCR amplification (Turner et al., 2014). The degradation of dissolved DNA can be mediated via multiple mechanisms, such as the formation of cross-linkages, hydrolysis of phosphodiester bonds, and thymine photodimerization that maintain the double-stranded structure of DNA detectable via fluorometry, but leave it unamplifiable with PCR (Bhoyar et al., 2024; Lindahl, 1993; Torti et al., 2015). Future studies should also consider the amplifiable quality of this dissolved DNA and use DNA repair procedures to make this dissolved DNA more usable (Bhoyar et al., 2024; Mouttham et al., 2015). Furthermore, the dissolved eDNA can also be removed from its dissolved form and get adsorbed to cells, suspended particles, or incorporated into microbial biofilms and not get isolated using the water filtration method in this study leading to a lower sequence richness (Brandão-Dias et al., 2023; Kirtane et al., 2023; Kirtane & Deiner, 2024; Power et al., 2023; Snyder et al., 2023).

The adsorbed eDNA state had the lowest DNA yield of the three states, but unlike the dissolved state, the adsorbed DNA was amplifiable in most samples, indicating that the DNA present in the adsorbed state might indeed be protected from degradation (Demaneche et al., 2001; Giguet-Covex et al., 2019; Paget et al., 1992). However, compared to the membrane-bound state, the adsorbed state did have a significantly lower sequence richness. The adsorbed eDNA might readily settle leading to its removal from the water column and accumulate in the sediment (Jerde et al., 2016; Shogren et al., 2017). Perhaps the adsorbed eDNA state would be better sampled directly from stream sediment instead of the water column, where it can increase the detection probability of rare species (Kirtane et al., 2019), or diversity of organisms (Sakata et al., 2020; Turner et al., 2015) despite the inability of being concentrated like filtration of water samples (Jerde et al., 2016; Shogren et al., 2017). Although physiochemical mechanisms could underlie the observed patterns, the lower yield and richness of adsorbed DNA may also result from methodological biases. Prior controlled experiments using the same workflow reported relatively low recovery of the adsorbed state (Kirtane et al., 2023). Notably, the taxonomic composition of the adsorbed state more closely resembled the membrane-bound state than the dissolved state, suggesting potential cross-over during the state-sorting process. This may be due in part to shared processing steps between the adsorbed and membrane-bound states, which could lead to contamination from lysed cells during transport or desorption. To enhance the separation of these fractions, a deeper understanding of adsorption mechanisms and membrane integrity is essential (Kirtane et al., 2023; Lever et al., 2015).

To our knowledge, this study is the first to investigate the diversity of eDNA in multiple states from environmental samples. Thus, the findings of this study cannot be generalized to all aquatic systems as factors such as temperature, discharge, pH, biological productivity, etc., can all influence the formation and persistence of eDNA states (Barnes & Turner, 2016; Eichmiller et al., 2016; Kirtane et al., 2023; Kirtane & Deiner, 2024; Seymour et al., 2018; Strickler et al., 2015). Even within this study, the sequence richness captured was influenced by the water chemistry. For instance, the two high-altitude Alpine watersheds, Sils and Silvaplana, exhibited notably higher richness in the dissolved DNA state compared to other watersheds and were characterized by lower temperature, pH, electrical conductivity, and dissolved solids (Figure S9, S13). State sorting studies need to be replicated in diverse environments where the factors impacting the formation and persistence of different states of eDNA can be systematically studied.

### Transport Dynamics of eDNA States in Lotic Systems

eDNA transport patterns were state-dependent, with the membrane-bound state showing a clear pattern: community similarity between flow-connected sites declined with increasing distance. This trend was observed for both ITS and COI markers, but only in the membrane-bound state. In contrast, many flow-connected samples in the dissolved and adsorbed states shared no ASVs, suggesting more rapid turnover—likely due to degradation (dissolved state) or settling (adsorbed state) (Figure 5, S12, S14). Although the distance-associated reduction in shared richness in the membrane-bound state was significant when all samples were pooled, it held true in only three of the eight individual watersheds (Figure S12, S14). While some other states and watersheds showed similar declines in shared richness with distance, these correlations were not statistically significant (Figure 5, S12).

Similar distance-associated declines in community similarity have been observed in small mountain streams (Reji Chacko et al., 2023), while studies of large river systems demonstrate that eDNA transport can extend over much greater distances, in some cases beyond 100 km (Perry et al., 2024). Here, we were limited by the length of the headwater streams to investigate transport beyond 18 km. Similarly, many studies reporting lower eDNA transport limits were unable to extend their sampling design to longer distances (Jo & Yamanaka, 2022). Studies reporting longer eDNA transport distances are more common in high-discharge rivers, where greater velocity and cross-sectional area increase downstream transport potential compared to lower-discharge headwater streams (Jo & Yamanaka, 2022). Downstream detection of eDNA also depends on its starting concentration, with higher starting concentrations at the source resulting in detectable eDNA further downstream after degradation, settling, and dilution (Sansom & Sassoubre, 2017). The concentrations of eDNA in larger rivers may be higher than in small headwater streams (Deiner et al., 2016; Perry et al., 2024). Furthermore, biofilms can increase eDNA removal rates, with smaller streams providing more biofilm contact and thereby enhancing removal (Rivera et al., 2023; Snyder et al., 2023). Future studies should characterize the effect of canal size and discharge on the transport potential of eDNA states.

### eDNA states at the inlet-lake interface

If rivers are hypothesized as ‘conveyor belts’ carrying biodiversity information (Deiner et al., 2016b), then lakes could potentially act as accumulators of this information, creating ‘hotspots’ for eDNA. Here, we find that the conveyor belt seems to be disrupted at the stream-lake confluence, as lake samples contained a largely different metazoan and plant community composition despite being collected only a short distance (2-50m) from the confluence. However, in some cases up to 60 % of the sequences found in the inlet samples were shared with the lake samples (Figure S15). For instance, eDNA from stream-dwelling species such as *Gammarus fossarum* and *Dinocras cephalotes* was also found in the lake samples, indicating eDNA transport from streams to lakes (Elbrecht et al., 2014; Pöckl et al., 2003). This suggests eDNA transport into the lakes; however, the exact fate of this transported eDNA remains unknown resulting in the lack of optimal sites for lake sampling. This turnover may not occur in lakes with low residence times or inlets with high discharge volumes. The observed shift in eDNA community composition at the stream–lake confluence may reflect underlying hydrodynamic processes, supporting the emerging view that eDNA can serve as a tracer for hydrological research (URycki et al., 2024).

The fate of river inflow within a lake is strongly influenced by how its density compares to that of the lake water. If the inflow is relatively buoyant, it tends to spread across the surface as an overflow, whereas denser inflows typically sink and follow the lakebed as underflows (Imberger & Patterson, 1989). Recent observations in Lake Geneva have revealed that the negatively buoyant Rhône mixing only after significant downstream travel into the metalimnion (Piton et al., 2022). In this study, we sampled the surface water of the lake, which was warmer than the streams (Figure S9). It may be that eDNA from the streams ends up in the metalimnion or the hypolimnion before dissipating into the lake (Littlefair et al., 2021). The fate of stream-borne eDNA in the lake remains uncertain, a question that invites further exploration beneath the surface.

This study highlights the need to consider eDNA states to investigate the physical processes governing eDNA transport in aquatic systems. ASV richness (COI marker) patterns varied by eDNA state: for membrane-bound DNA, stream and inlet samples showed significantly higher richness than lake samples, whereas, in the dissolved state, richness was greater in lake samples than in streams or inlets (Figure S15). In lotic systems like streams, turbulent flow may keep cells and adsorbents suspended, whereas the lentic conditions in lakes may promote their sedimentation, reducing their presence in surface waters or transport into the interiors of the lake (Biernaski et al., 2025; Pont, 2024; Wang et al., 2023; Xiong et al., 2025). Dissolved state on the other hand, would not be susceptible to this settling. Despite this evidence, there was still a strong change in community composition at the stream-lake interface in the dissolved state, suggesting that the settling behavior might not be the only factor responsible for the observed phenomena.

### Limitations and outlook

This study design’s main limitation with empirically studying eDNA transport is the uncertainty of whether the eDNA captured at a downstream site originated from an upstream source or was released locally. To negate the influence of new eDNA released between the upstream and downstream sites, using a fixed source of eDNA input such as fish farms, caged fish, or spiked non-native eDNA in a stream can be incorporated to resolve this limitation (Jane et al., 2015; Nukazawa et al., 2018; Perry et al., 2024). Combining species occupancy modeling can help identify patchily distributed species in the landscape suitable for empirically investigating the transport behavior of eDNA without experimental eDNA spikes (Brantschen et al., 2024; Burian et al., 2021).

Another limitation of this study is the collection of one sample replicate per site. Past studies have shown eDNA signals are heterogeneous and impacted by pulse inputs of eDNA, season, rainfall, diurnal/nocturnal activity, etc. (Shiozuka et al., 2023; Suzuki et al., 2022; Yang et al., 2021). The state sorting methodology also limited the sample volume to 100 mL, lower than most typically sampled 1000 mL which has been shown to capture higher species diversity (Sakata et al., 2021). As this was among the first studies of its kind, our lake sampling was limited to surface waters, leaving uncertainty about where stream eDNA ultimately settles. A more refined design, involving sampling across depths and at varying distances from the inlet, could reveal whether stream-derived eDNA is concentrated in deeper strata or removed from the water column via sedimentation. Despite these constraints, our findings highlight the value of separating eDNA into its states for revealing underlying transport processes and community patterns, reinforcing the potential of this approach for improving catchment-scale biodiversity assessments.

## Acknowledgments

We thank Dr. Noriko Uchida for their help in collecting field samples. We thank Dr. Catia Lucio Pereira for helping design custom sequencing primers and universal primers with MIDs. We thank Dr. Jean-Claude Walser and Dr. Cátia Lúcio Pereira for help with bioinformatics. Data produced and analyzed in this paper were generated in collaboration with the Genetic Diversity Centre (GDC), ETH Zurich and the Functional Genomics Center Zurich (FGCZ), University of Zurich.

## Author Contributions

AK and KD conceptualized the project. AK, KD, and EL performed field sampling. AK, ZD, and EL performed laboratory analysis. Laboratory analysis of the dissolved eDNA state was performed as part of a master’s thesis by ZD. All authors contributed to the data analysis. AK wrote the first draft. All authors contributed to writing and editing the final draft.

