## Supplementary material for "Sorting States of Environmental DNA Reveals State-Specific Biodiversity Signals and Transport Patterns in Eight Alpine Watersheds": text

**Supporting information**

Table S1: The number of streams sampled, the number of samples collected from each Lake watershed, with the average pH, temperature, electrical conductivity, and total dissolved solids of the sampled water.

| Lake watershed | Number of streams sampled | Number of samples collected | Temperature (°C) | pH | TDS (ppm) | EC (mS/cm) |
|---|---|---|---|---|---|---|
| Aegeri | 6 | 23 | 14.44 ± 1.76 | 8.03 ± 0.24 | 250.00 ± 31.38 | 0.35 ± 0.05 |
| Baldegg | 9 | 42 | 18.66 ± 2.58 | 7.99 ± 0.31 | 355.76 ± 56.62 | 0.48 ± 0.08 |
| Greifen | 7 | 42 | 19.96 ± 2.50 | 7.95 ± 0.33 | 394.06 ± 83.50 | 0.55 ± 0.12 |
| Hallwil | 7 | 27 | 17.27 ± 2.43 | 8.15 ± 0.27 | 344.90 ± 95.86 | 0.47 ± 0.13 |
| Lauerz | 7 | 31 | 14.96 ± 1.72 | 8.03 ± 0.20 | 225.58 ± 36.58 | 0.31 ± 0.05 |
| Pfäffikon | 1 | 10 | 10.20 ± 3.50 | 8.17 ± 0.19 | 319.86 ± 62.52 | 0.43 ± 0.12 |
| Sils | 4 | 18 | 10.95 ± 2.64 | 7.81 ± 0.28 | 110.29 ± 42.80 | 0.15 ± 0.06 |
| Silvaplana | 6 | 28 | 11.12 ± 3.14 | 7.93 ± 0.31 | 84.33 ± 44.28 | 0.11 ± 0.04 |

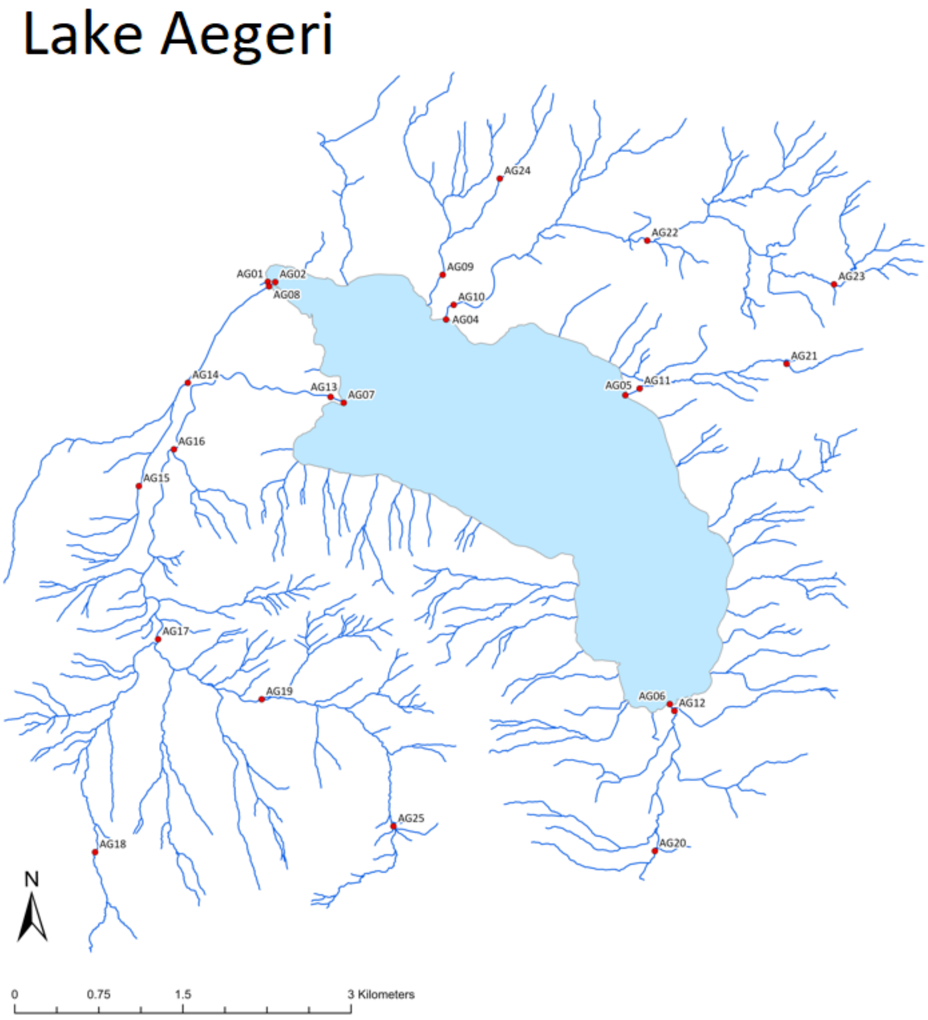


Figure S1: Map of Lake Aegeri watershed showing sampling locations indicated by red points.

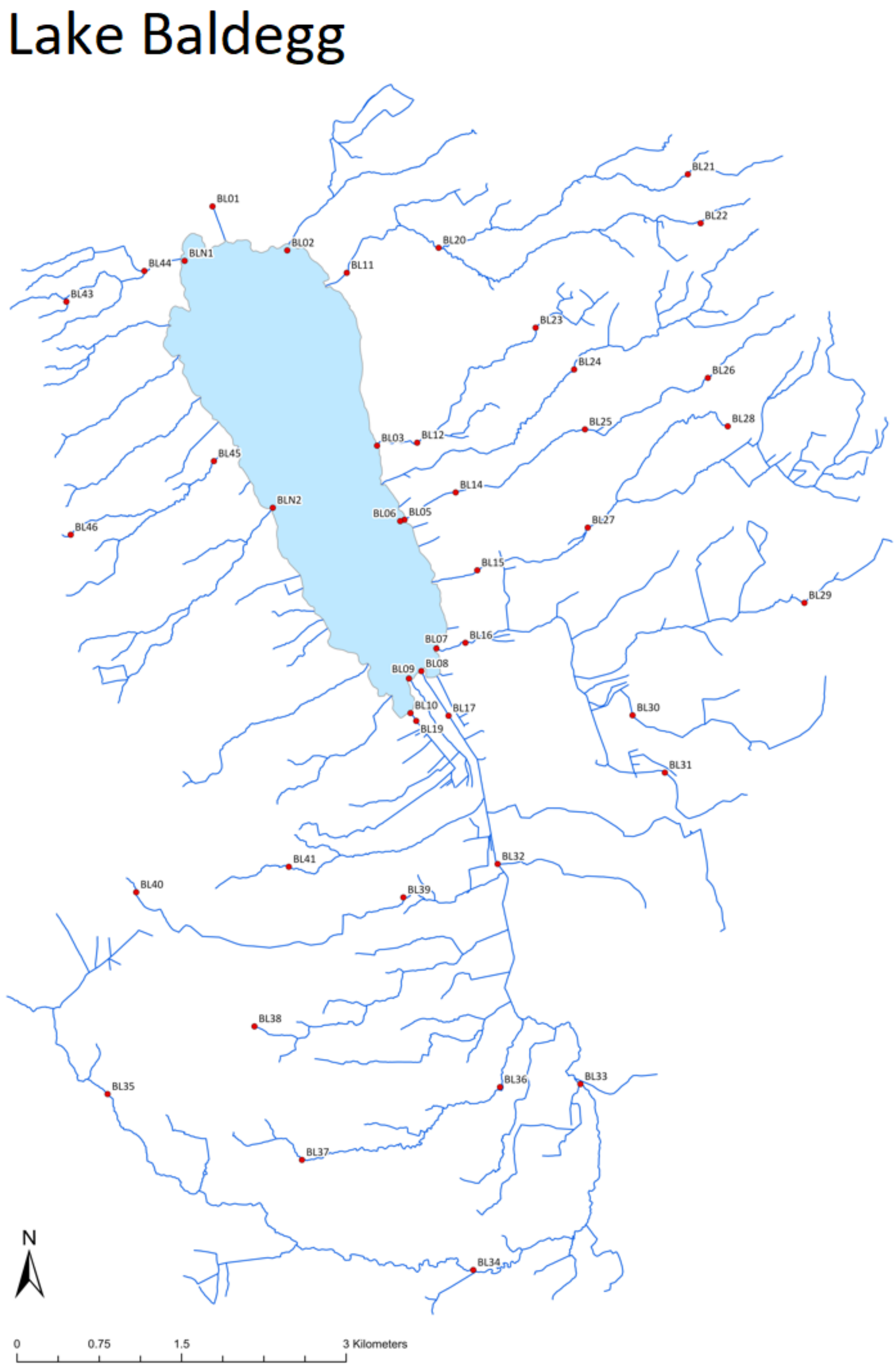


Figure S2: Map of Lake Baldegg watershed showing sampling locations indicated by red points.

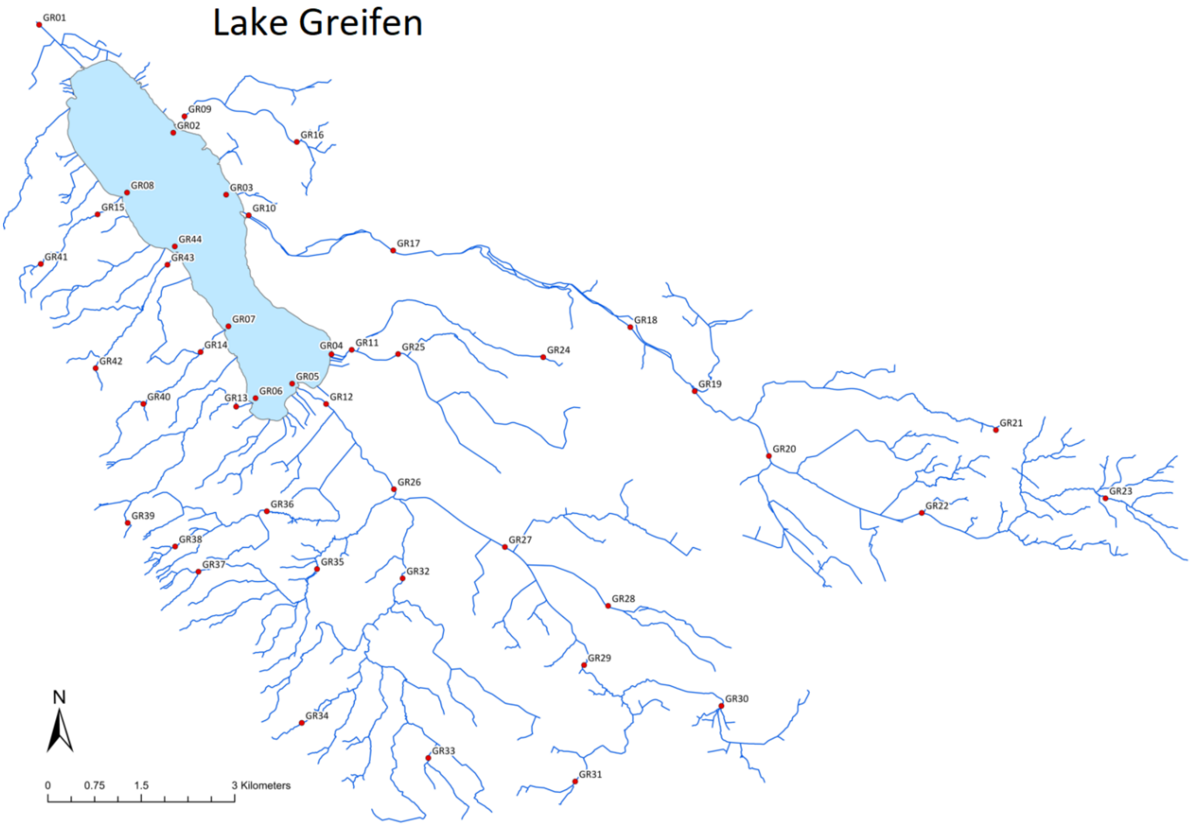


Figure S3: Map of Lake Greifen watershed showing sampling locations indicated by red points.

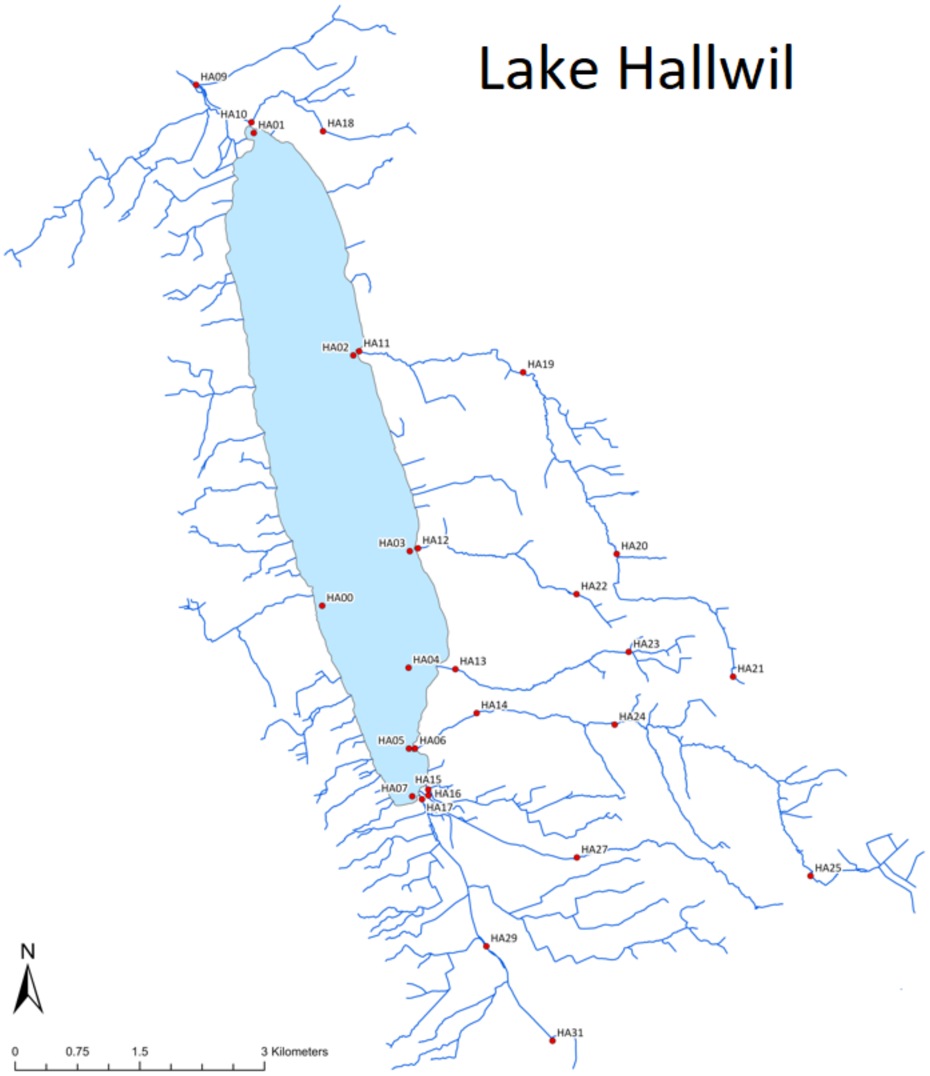


Figure S4: Map of Lake Hallwil watershed showing sampling locations indicated by red points.

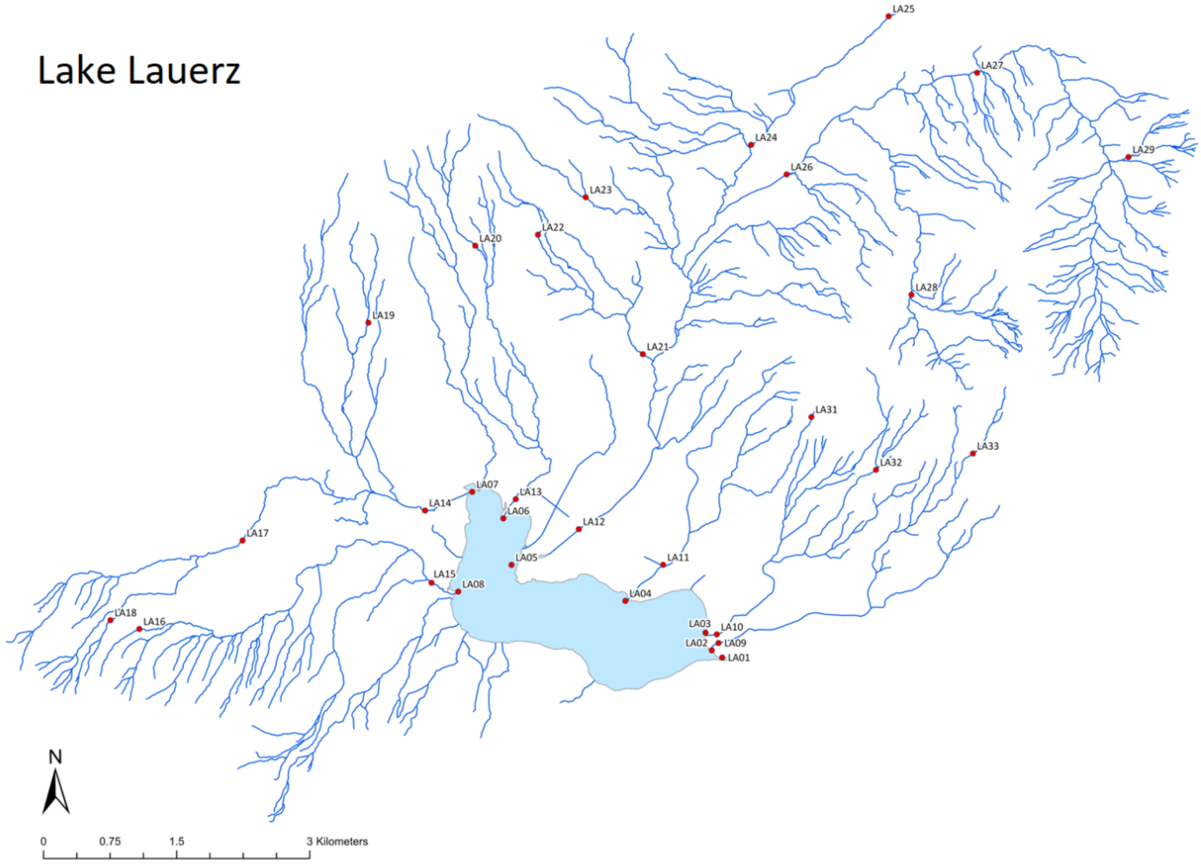


Figure S5: Map of Lake Lauerz watershed showing sampling locations indicated by red points.

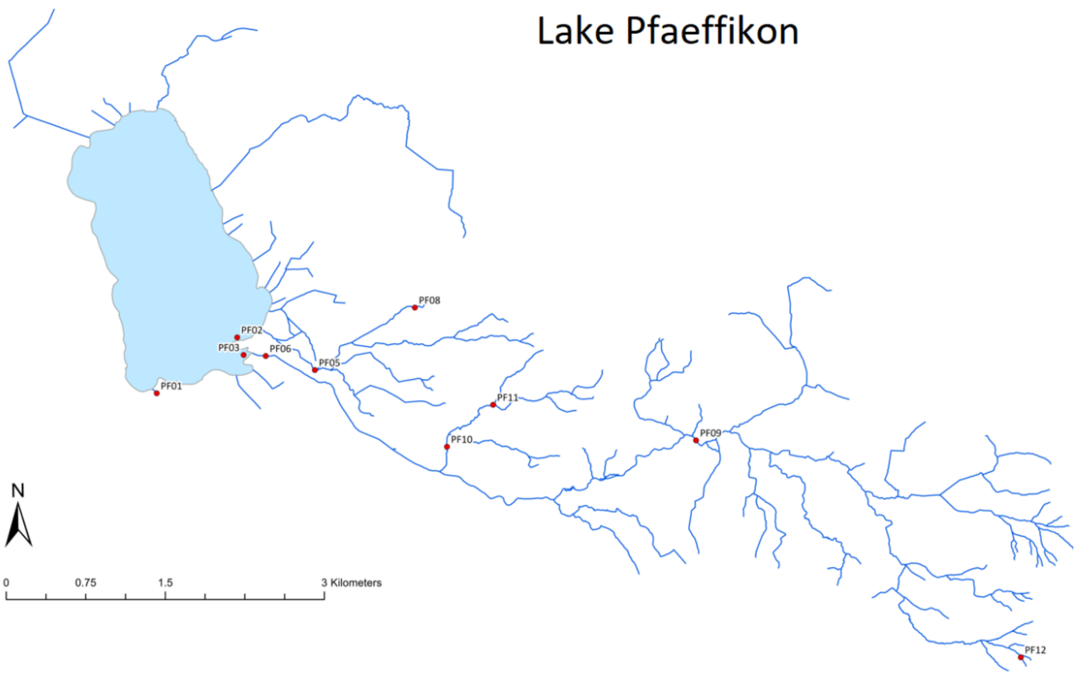


Figure S6: Map of Lake Pfaeffikon watershed showing sampling locations indicated by red points.

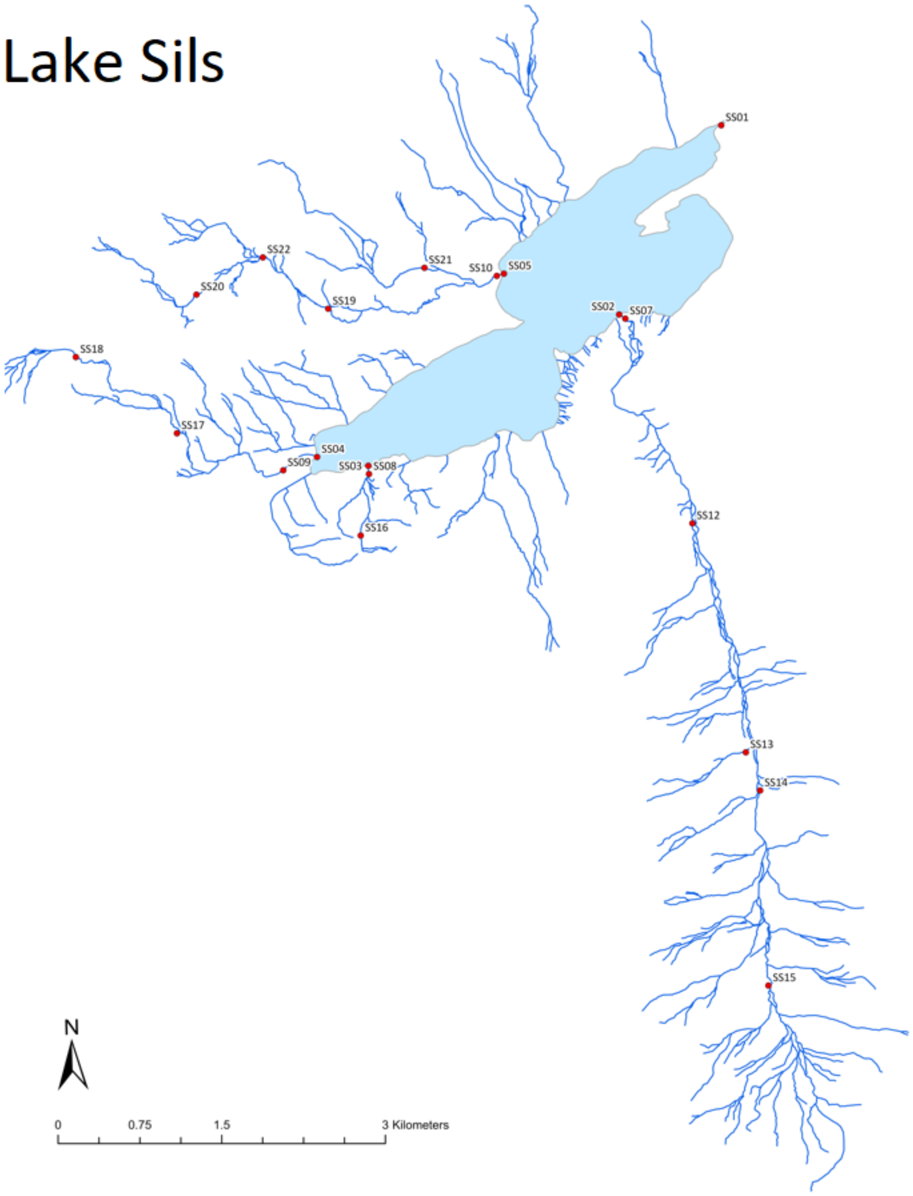


Figure S7: Map of Lake Sils watershed showing sampling locations indicated by red points.

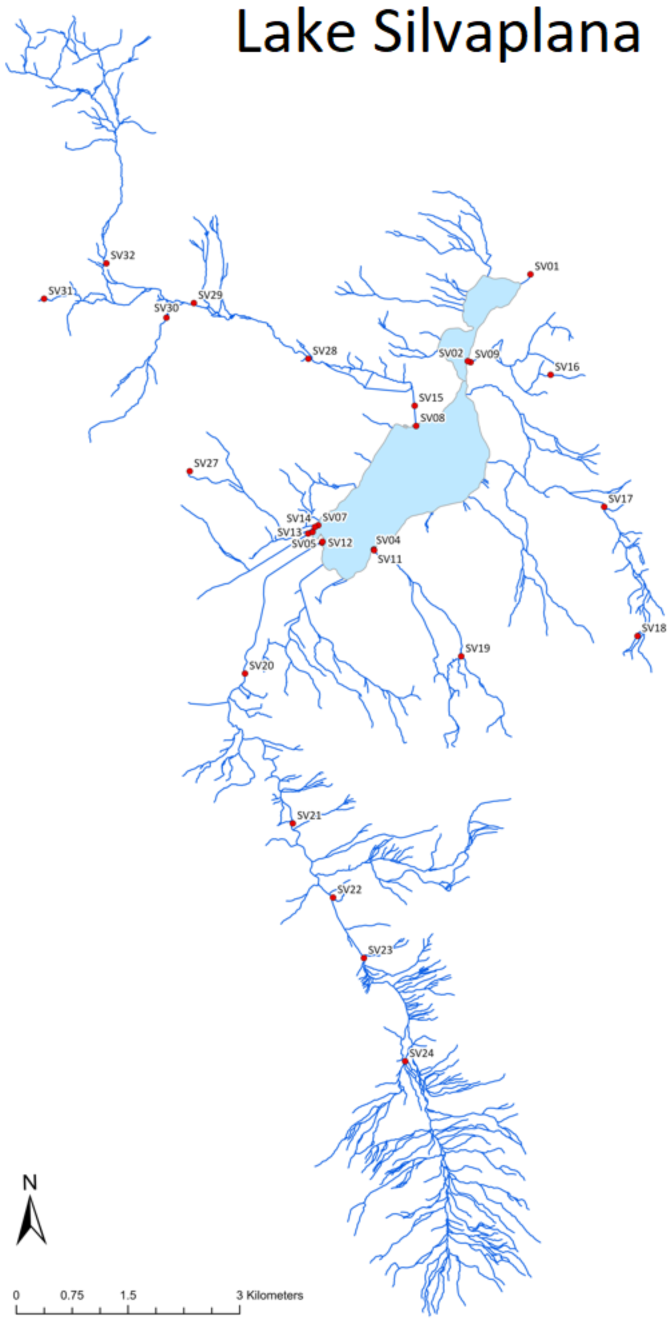


Figure S8: Map of Lake Silvaplana watershed showing sampling locations indicated by red points.

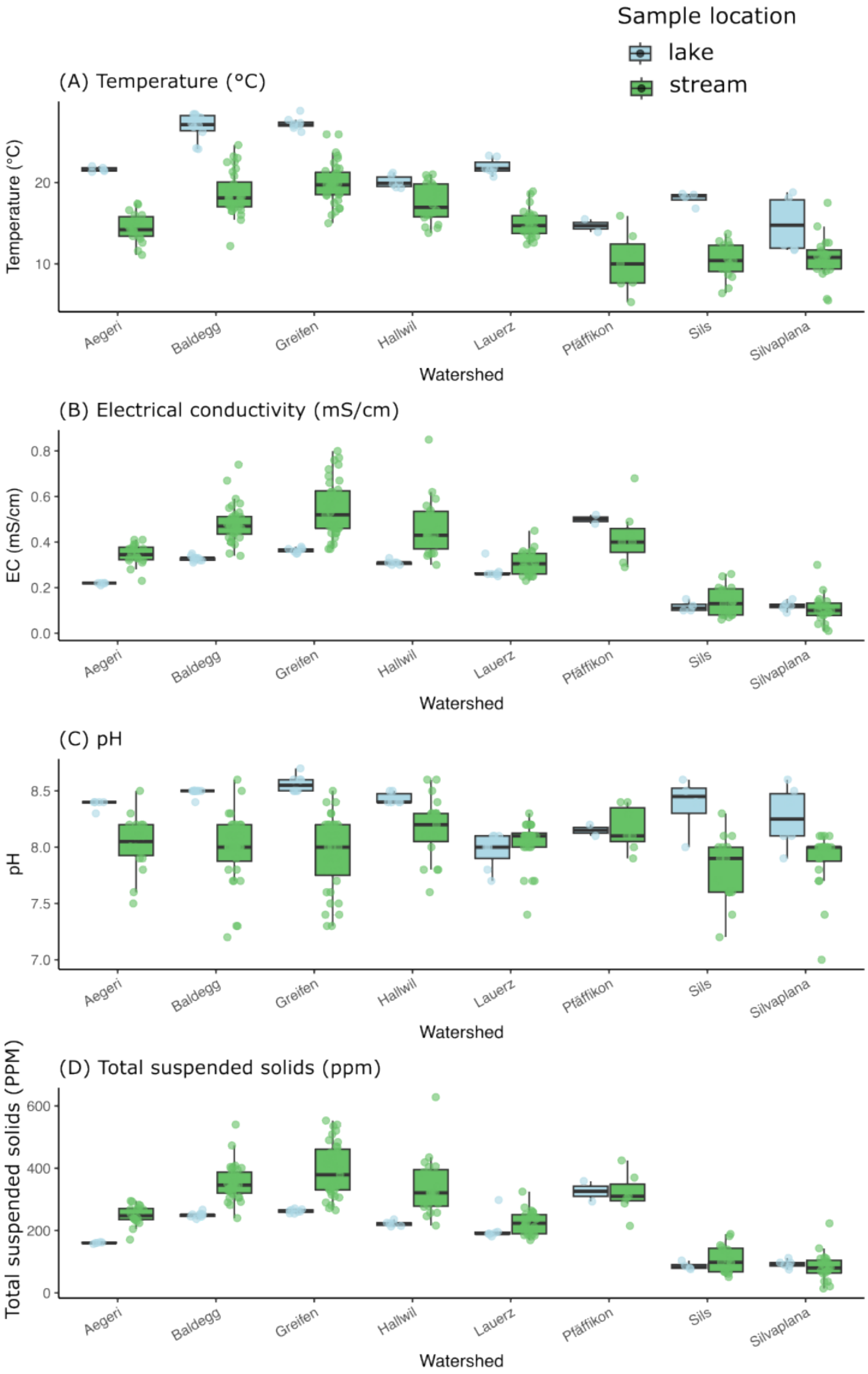


Figure S9: Metadata for (A) temperature, (B) electrical conductivity, (C) pH, and (D) total dissolved solids collected alongside water samples.

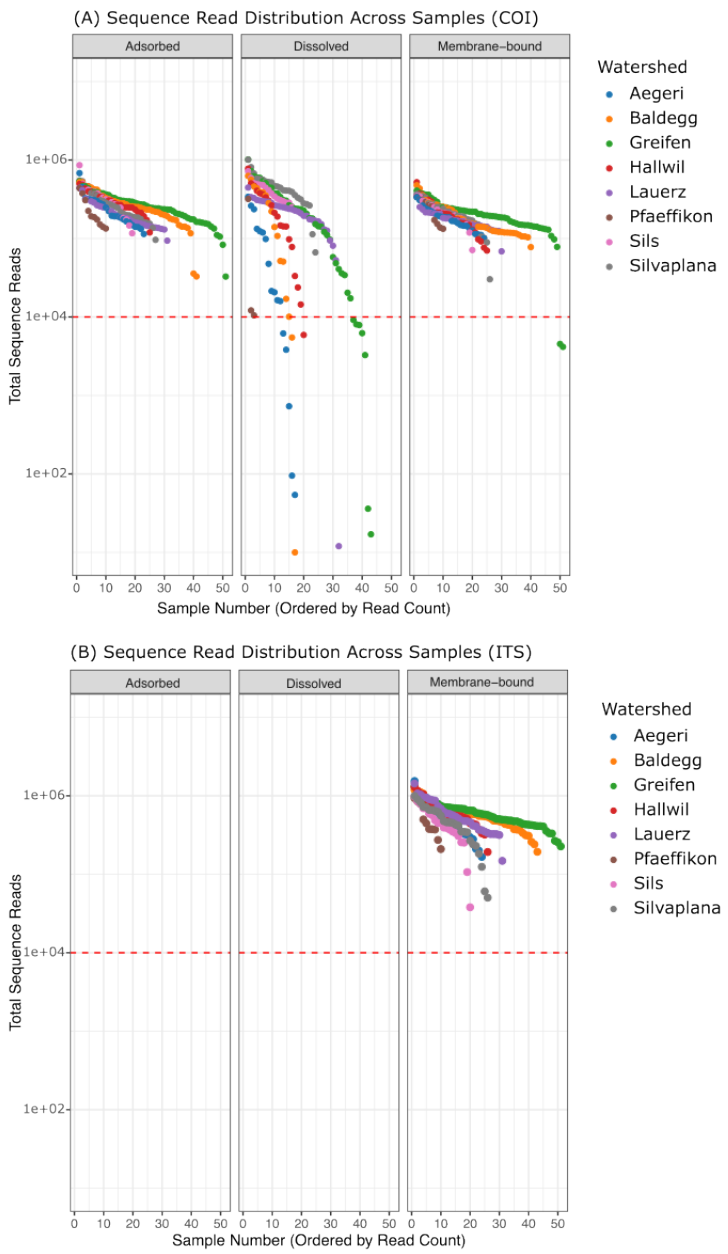


Figure S10: Sequence read counts for (A) COI and (B) ITS samples before rarefaction. The red horizontal line is drawn at the 10,000 reads threshold used to rarify the samples. All samples below the line were excluded from all analyses.

Table S2: PCR primers used in this study. Underlined letters indicate short tails, and bold letters indicate the location of the Illumina adaptors (P5 and P7). Brackets indicate the position of individual MIDs, which differ for each sample (Appendix Table 5). All primers were obtained from Integrated DNA Technologies, Coralville, Iowa.

| **Primer name** | **Sequence** | **Type** | **Role** | **Reference** |
|---|---|---|---|---|
| *Sauron-S878* | GGDRCWGGWTGAACWGTWTAYCCNCC | Forward | Boost primer | Rennstam Rubbmark et al. 2018 |
| *HCO2198* | TAAACTTCAGGGTGACCAAAAAATCA | Reverse | Boost primer | Folmer et al. 1994 |

| | | | | |
|---|---|---|---|---|
| *Sauron-S878-Tail* | CACCTGCTTCTAAATGGDRCWGGWTGAACWGTWTAYCCNCC | Forward | Universal primer with tail 1 | Rennstam Rubbmark et al. 2018 |
| *HCO2198-Tail* | CACTTCGACTCTTTACTAAACTTCAGGGTGACCAAAAAATCA | Reverse | Universal primer with tail 1 | Rennstam Rubbmark et al. 2018 |
| *P5-i5-Tail* | **AATGATACGGCGACCACCGAGATCTACAC**[insert individual MID per sample]CACCTGCTTCTAAAT | Forward | Index primer: Illumina sequencing adapters, MIDs, and tail 2 | (Rennstam Rubbmark et al. 2018) |
| *P7-i7 -Tail* | **CAAGCAGAAGACGGCATACGAGAT**[insert individual MID per sample]CACTTCGACTCTTTAC | Reverse | Index primer: Illumina sequencing adapters, MIDs, and tail 2 | (Rennstam Rubbmark et al. 2018) |
| *SEQ-Sauron* | CACCTGCTTCTAAATGGDRCWGGWTGAACWGTWTAYCCNCC | | Sequencing Primer COI | (Rennstam Rubbmark et al. 2018) |
| *IND-Sauron* | GGNGGRTAWACWGTTCAWCCWGYHCCATTTAGAAGCAGGTG | Reverse complement of SEQ-Sauron | Sequencing Primer COI | (Rennstam Rubbmark et al. 2018) |
| *SEQ-HCO2198* | CACTTCGACTCTTTACTAAACTTCAGGGTGACCAAAAAATCA | | Sequencing Primer COI | (Rennstam Rubbmark et al. 2018) |
| *IND-HCO2198* | TGATTTTTTGGTCACCCTGAAGTTTAGTAAAGAGTCGAAGTG | Reverse complement of SEQ-HCO2198 | Sequencing Primer COI | (Rennstam Rubbmark et al. 2018) |
| *ITS2F* | ATGCGATACTTGGTGTGAAT | Forward | Boost Primer | (Chen et al., 2010) |
| *ITS2R* | GACGCTTCTCCAGACTACAAT | Reverse | Boost Primer | (Chen et al., 2010) |
| *ITS2F-Tail* | CACCTGCTTCTAAATATGCGATACTTGGTGTGAAT | Forward | Universal primer with tail 1 | This study |
| *ITS2R-Tail* | CACTTCGACTCTTTACGACGCTTCTCCAGACTACAAT | Reverse | Universal primer with tail 1 | This study |
| *SEQ-ITS2F* | CACCTGCTTCTAAATATGCGATACTTGGTGTGAAT | | Sequencing Primer ITS2 | This study |
| *IND-ITS2F* | ATTCACACCAAGTATCGCATATTTAGAAGCAGGTG | reverse complement of SEQ-ITS2F | Sequencing Primer ITS2 | This study |
| *SEQ-ITS3R* | CACTTCGACTCTTTACGACGCTTCTCCAGACTACAAT | | Sequencing Primer ITS2 | This study |
| *IND-ITS3R* | ATTGTAGTCTGGAGAAGCGTCGTAAAGAGTCGAAGTG | reverse complement of SEQ-ITS3R | Sequencing Primer ITS2 | This study |
| *P5 Illumina adapter* | **AATGATACGGCGACCACCGA** | Forward | Reconditioning PCR | Illumina, San Diego |
| *P7 Illumina adapter* | **CAAGCAGAAGACGGCATACGA** | Reverse | Reconditioning PCR | Illumina, San Diego |

Table S3: Spearman correlation analysis based on the percent shared ASV richness and SUSO difference for each marker, state, and lake watershed combinations.

| Marker | Lake | State | n_pairs | Spearman's rho | p_value |
|---|---|---|---|---|---|
| COI | Agaeri | ADS | 22 | -0.4040059 | 0.06221757 |

| | | | | | |
|---|---|---|---|---|---|
| COI | Agaeri | DIS | 6 | 0.36742346 | 0.47366589 |
| COI | Agaeri | MEM | 22 | -0.5998985 | 0.00316469 |
| COI | Baldegg | ADS | 37 | -0.1915387 | 0.25610348 |
| COI | Baldegg | DIS | 1 | NA | NA |
| COI | Baldegg | MEM | 34 | -0.6168523 | 0.00010222 |
| COI | Greifen | ADS | 59 | -0.2621454 | 0.04488602 |
| COI | Greifen | DIS | 24 | -0.3500819 | 0.09353492 |
| COI | Greifen | MEM | 55 | -0.371271 | 0.00525971 |
| COI | Hallwil | ADS | 13 | -0.4306269 | 0.14184831 |
| COI | Hallwil | DIS | 7 | NA | NA |
| COI | Hallwil | MEM | 15 | -0.3941184 | 0.14605556 |
| COI | Lauerz | ADS | 22 | 0.03999324 | 0.85973986 |
| COI | Lauerz | DIS | 28 | -0.0724684 | 0.71401737 |
| COI | Lauerz | MEM | 27 | -0.0505199 | 0.80239629 |
| COI | Phaeffikon | ADS | 7 | -0.3162278 | 0.48958974 |
| COI | Phaeffikon | DIS | 1 | NA | NA |
| COI | Phaeffikon | MEM | 7 | -0.4743417 | 0.28217986 |
| COI | Sils | ADS | 17 | 0.0115239 | 0.96498716 |
| COI | Sils | DIS | 11 | 0.0978232 | 0.77477162 |
| COI | Sils | MEM | 21 | -0.0541277 | 0.8157411 |
| COI | Silvaplana | ADS | 26 | 0.18497802 | 0.36564523 |
| COI | Silvaplana | DIS | 21 | -0.3557727 | 0.1134625 |
| COI | Silvaplana | MEM | 26 | -0.0055647 | 0.97847649 |
| ITS | Agaeri | MEM | 22 | -0.5192504 | 0.01326927 |
| ITS | Baldegg | MEM | 38 | -0.1840464 | 0.26867442 |
| ITS | Greifen | MEM | 57 | -0.3569243 | 0.00642195 |
| ITS | Hallwil | MEM | 15 | -0.4436524 | 0.09762019 |
| ITS | Lauerz | MEM | 30 | -0.2717512 | 0.14630773 |
| ITS | Phaeffikon | MEM | 7 | -0.4743417 | 0.28217986 |
| ITS | Sils | MEM | 21 | -0.4064358 | 0.06750084 |
| ITS | Silvaplana | MEM | 26 | 0.10425281 | 0.61227566 |

**Table S4: P***airwise PERMANOVA comparisons of community composition between lake positions (stream, inlet, lake, outflow) across all three states for COI and membrane-bound state for ITS. Values shown are F-statistics (F_model), $R^2$, unadjusted p-values, and Bonferroni-adjusted p-values (p_adj) based on Jaccard distance and 999 permutations. (DIS = Dissolved, MEM = Membrane-bound, ADS = Adsorbed)*

| Dataset | Group1 | Group2 | F_model | R2 | p_value | p_adj |
|---|---|---|---|---|---|---|
| DIS_COI | stream | inlet | 1.02 | 0.0109 | 0.402 | 1 |
| DIS_COI | inlet | lake | 2.52 | 0.047 | 0.001 | 0.024 |
| DIS_COI | lake | outflow | 0.618 | 0.0149 | 0.981 | 1 |

| | | | | | | |
|---|---|---|---|---|---|---|
| DIS_COI | stream | lake | 3.46 | 0.0316 | 0.001 | 0.024 |
| DIS_COI | outflow | inlet | 1.71 | 0.0752 | 0.003 | 0.072 |
| DIS_COI | outflow | stream | 1.41 | 0.018 | 0.011 | 0.264 |
| MEM_COI | stream | inlet | 1.17 | 0.00729 | 0.026 | 0.624 |
| MEM_COI | inlet | lake | 4.73 | 0.0505 | 0.001 | 0.024 |
| MEM_COI | lake | outflow | 0.652 | 0.0124 | 0.961 | 1 |
| MEM_COI | stream | lake | 6.15 | 0.037 | 0.001 | 0.024 |
| MEM_COI | outflow | inlet | 2.13 | 0.04 | 0.001 | 0.024 |
| MEM_COI | outflow | stream | 2.1 | 0.0169 | 0.001 | 0.024 |
| ADS_COI | stream | inlet | 1.06 | 0.00659 | 0.138 | 1 |
| ADS_COI | inlet | lake | 4.33 | 0.0454 | 0.001 | 0.024 |
| ADS_COI | lake | outflow | 0.667 | 0.0124 | 0.935 | 1 |
| ADS_COI | stream | lake | 5.55 | 0.0331 | 0.001 | 0.024 |
| ADS_COI | outflow | inlet | 1.97 | 0.0366 | 0.001 | 0.024 |
| ADS_COI | outflow | stream | 1.93 | 0.0154 | 0.001 | 0.024 |
| MEM_ITS | stream | inlet | 1.16 | 0.00692 | 0.112 | 1 |
| MEM_ITS | inlet | lake | 4.51 | 0.0482 | 0.001 | 0.024 |
| MEM_ITS | lake | outflow | 0.789 | 0.0152 | 0.877 | 1 |
| MEM_ITS | stream | lake | 5.27 | 0.0312 | 0.001 | 0.024 |
| MEM_ITS | outflow | inlet | 1.79 | 0.0333 | 0.002 | 0.048 |
| MEM_ITS | outflow | stream | 1.72 | 0.0134 | 0.003 | 0.072 |

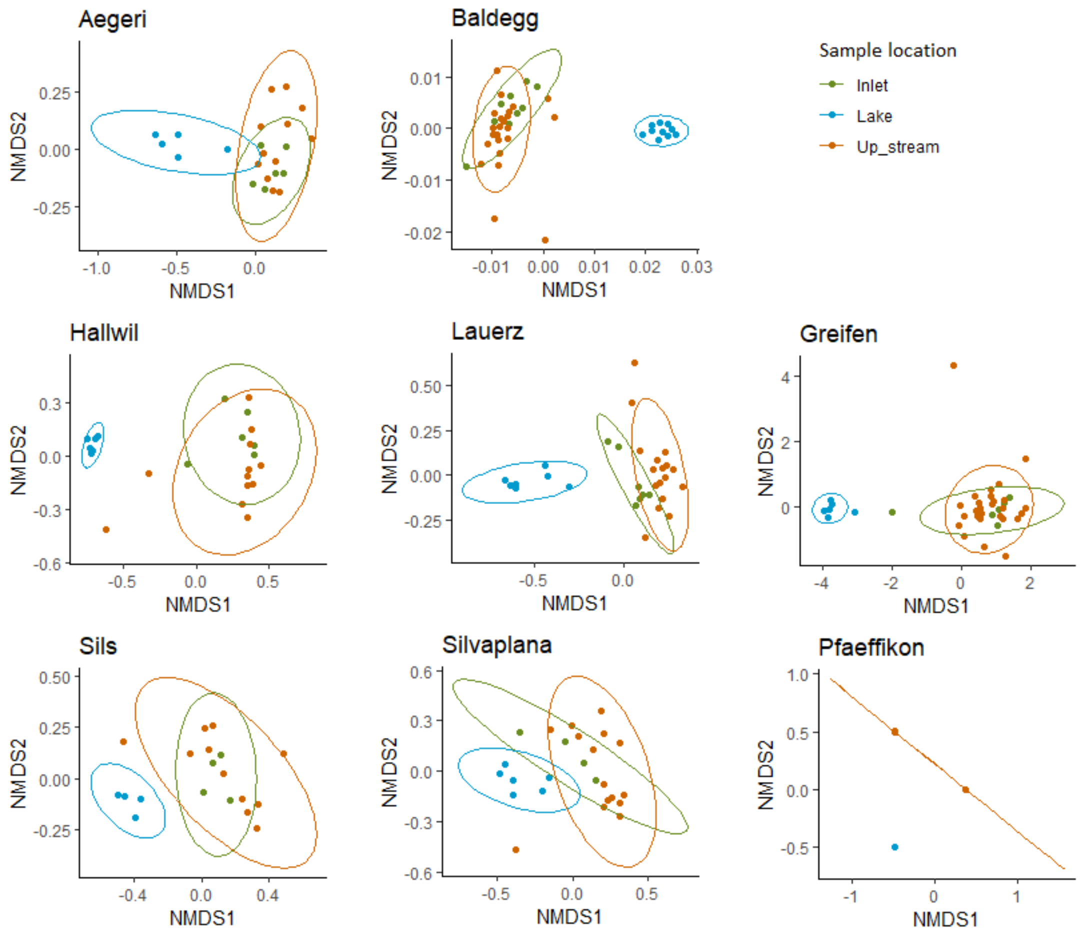


Figure S11. NMDS plots with the COI community composition across sampling locations with combined data from all three states. Only samples where all three states were sequenced successfully are used in this figure. The ellipses represent 95 % confidence intervals for COI communities captured from up-stream, inlet, and lake samples. Lake Greifen and Hallwil were sampled 50 m from the inlet, while other lakes were sampled closer to the inlet.

Table S5: Pairwise PERMANOVA results for stream, inlet, and lake samples for each of the eight lakes. Inlet samples from Lake Pfaeffikon were excluded from the analysis due to insufficient sample size. Data from all three states were combined for this analysis.

| **Lake** | **pairs** | **Df** | **SumsOfSqs** | **F.Model** | **$R^2$** | **p.value** | **p.adjusted** | **sig** |
|---|---|---|---|---|---|---|---|---|
| Aegeri | Lake vs Inlet | 1 | 1.1569892 | 3.2693483 | 0.26646471 | 0.003 | 0.009 | * |
| | Lake vs stream | 1 | 1.3580598 | 3.3797111 | 0.18388271 | 0.001 | 0.003 | * |
| | Inlet vs stream | 1 | 0.4583195 | 0.9783158 | 0.05762148 | 0.588 | 1 | |
| Baldegg | Lake vs Inlet | 1 | 1.6028464 | 4.4127315 | 0.2060798 | 0.001 | 0.003 | * |
| | Lake vs stream | 1 | 2.1917043 | 5.3786931 | 0.14785284 | 0.001 | 0.003 | * |
| | Inlet vs stream | 1 | 0.4841829 | 1.0273685 | 0.03311169 | 0.284 | 0.852 | |
| Greifen | Lake vs Inlet | 1 | 1.4350613 | 4.3740384 | 0.26713254 | 0.001 | 0.003 | * |
| | Lake vs stream | 1 | 2.0834232 | 4.9422139 | 0.13378229 | 0.001 | 0.003 | * |
| | Inlet vs stream | 1 | 0.4856319 | 1.030732 | 0.03120524 | 0.259 | 0.777 | |
| Hallwil | Lake vs stream | 1 | 1.3202623 | 3.2075749 | 0.16699531 | 0.001 | 0.003 | * |
| | Lake vs Inlet | 1 | 1.1213352 | 3.0214748 | 0.23203784 | 0.003 | 0.009 | * |
| | stream vs Inlet | 1 | 0.4831775 | 1.0283774 | 0.06039198 | 0.277 | 0.831 | |
| Lauerz | Lake vs Inlet | 1 | 1.0329995 | 2.5404951 | 0.17471861 | 0.002 | 0.006 | * |
| | Lake vs stream | 1 | 1.2320637 | 2.8123488 | 0.11334472 | 0.001 | 0.003 | * |
| | Inlet vs stream | 1 | 0.572178 | 1.2301017 | 0.05295292 | 0.001 | 0.003 | * |
| Pfaeffikon | Lake vs Inlet | | | | | | | |
| | Lake vs stream | 1 | 0.7360524 | 1.7987724 | 0.26457312 | 0.044 | 0.132 | |
| | Inlet vs stream | | | | | | | |
| Sils | Lake vs Inlet | 1 | 0.9714021 | 2.9204381 | 0.32738729 | 0.039 | 0.117 | |
| | Lake vs stream | 1 | 1.1259216 | 2.730764 | 0.17359385 | 0.001 | 0.003 | * |
| | Inlet vs stream | 1 | 0.5445968 | 1.1774646 | 0.08305184 | 0.068 | 0.204 | |
| Silvaplana | Lake vs Inlet | 1 | 0.7329086 | 1.7952526 | 0.18327782 | 0.01 | 0.03 | . |
| | Lake vs stream | 1 | 1.0310707 | 2.3386669 | 0.10959761 | 0.001 | 0.003 | * |
| | Inlet vs stream | 1 | 0.4880291 | 1.0386392 | 0.05757858 | 0.258 | 0.774 | |

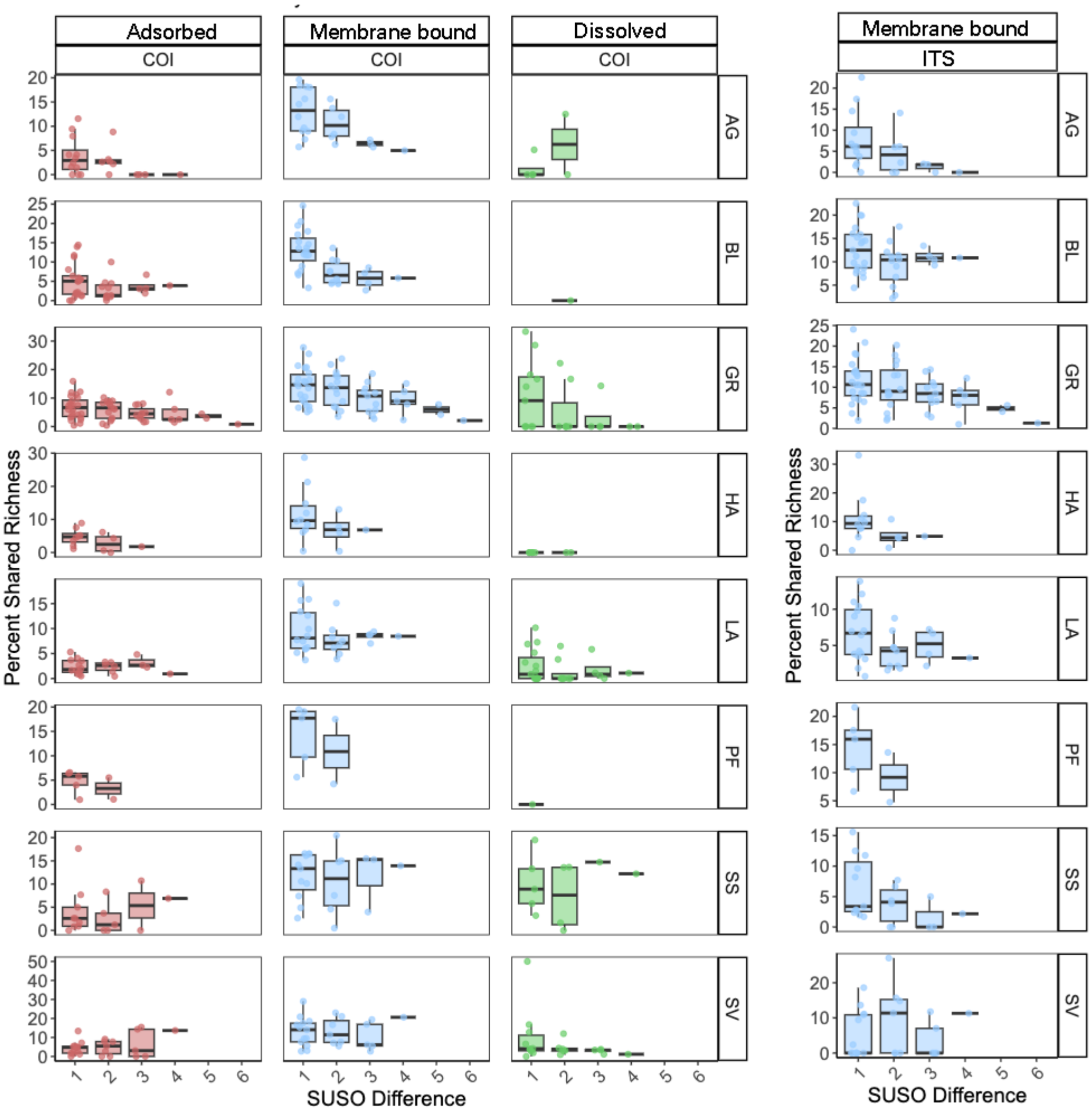


Figure S12: Percentage of shared species richness over SUSO between flow-connected sample pairs for adsorbed, membrane-bound, and dissolved states for COI and ITS markers for streams in each of the eight lake watersheds investigated.

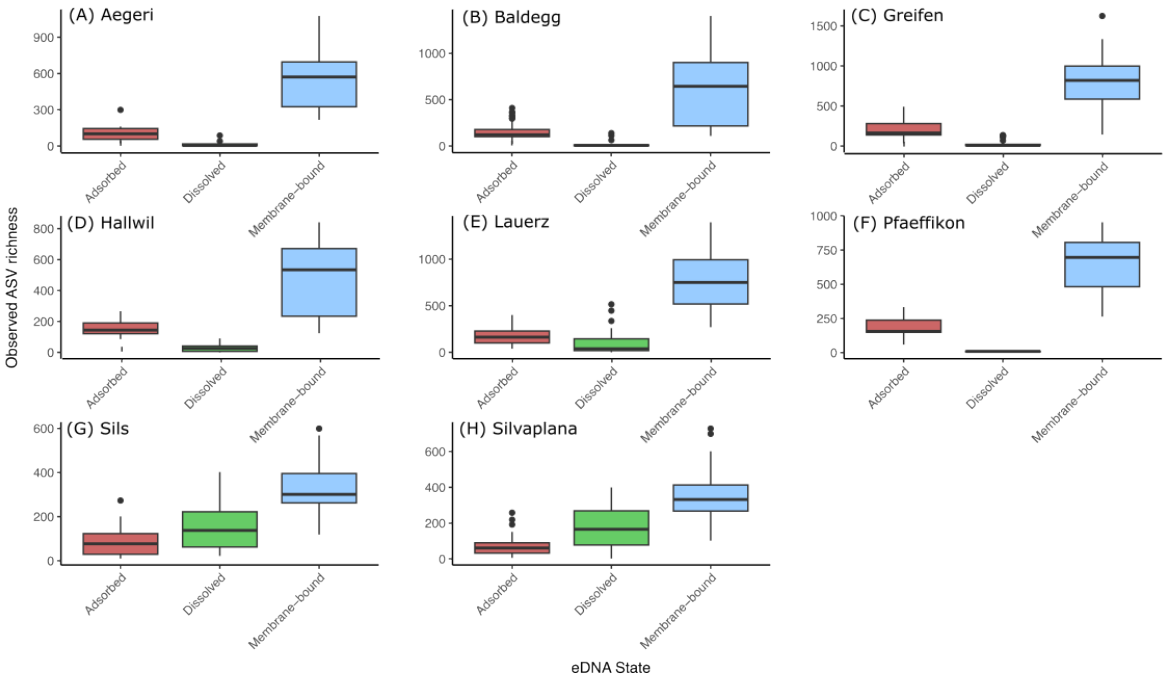


Figure S13: Boxplot of ASV richness from stream samples in adsorbed, dissolved, and membrane-bound eDNA in each of the eight lake watersheds sampled.

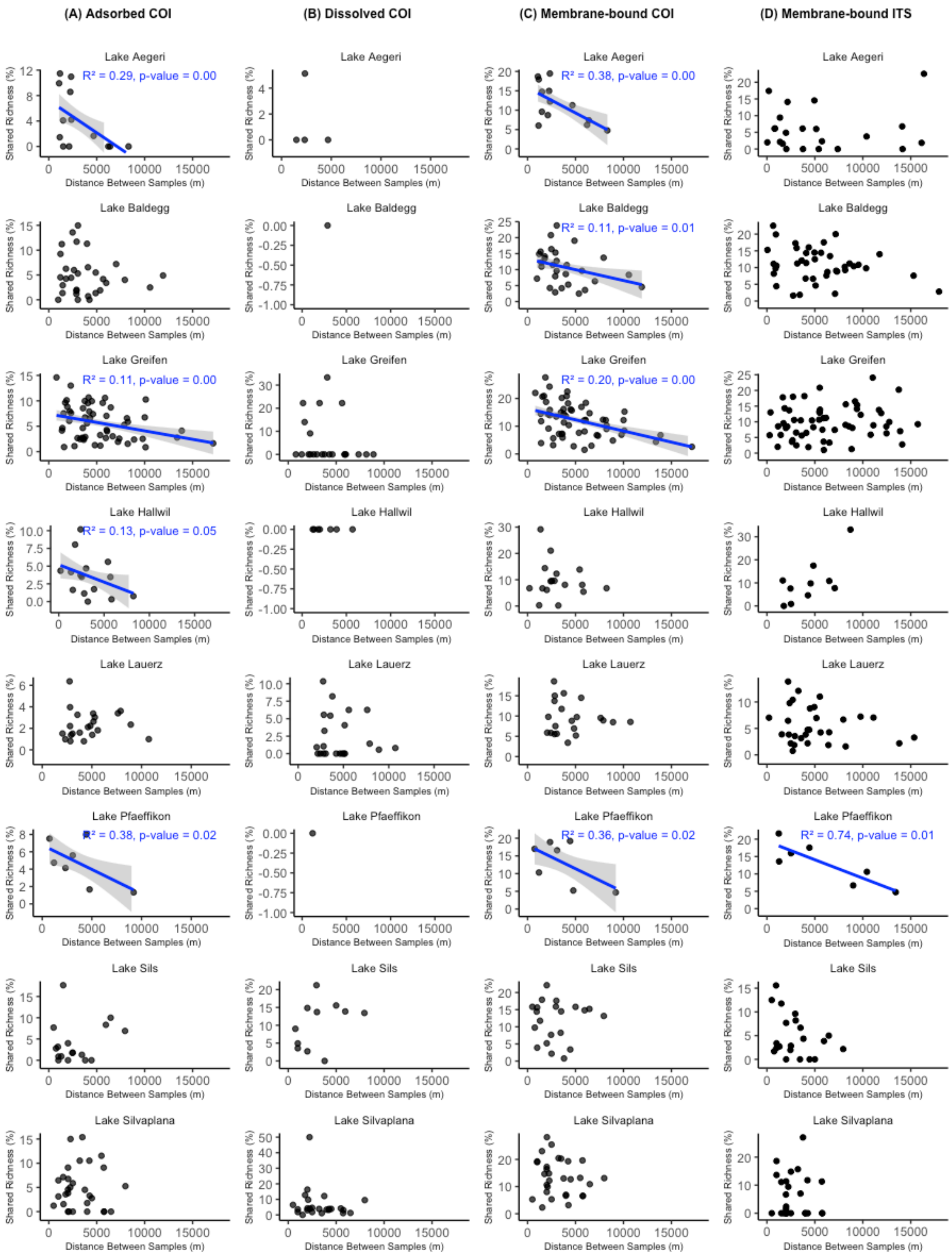


Figure S14. Percentage of shared ASV richness over distance between flow-connected sample pairs for (A) Adsorbed, (B) Membrane-bound, and (C) Dissolved states of eDNA for the COI marker and (D) Membrane-bound DNA for the ITS marker for streams in each of the eight lake watersheds investigated. The blue line indicates the linear regression line with the shared area showing a 95 % confidence interval. R-squared and p values of the regression fit are displayed in each plot. Plots without trendlines have a non-significant relationship based on the linear regression analysis.

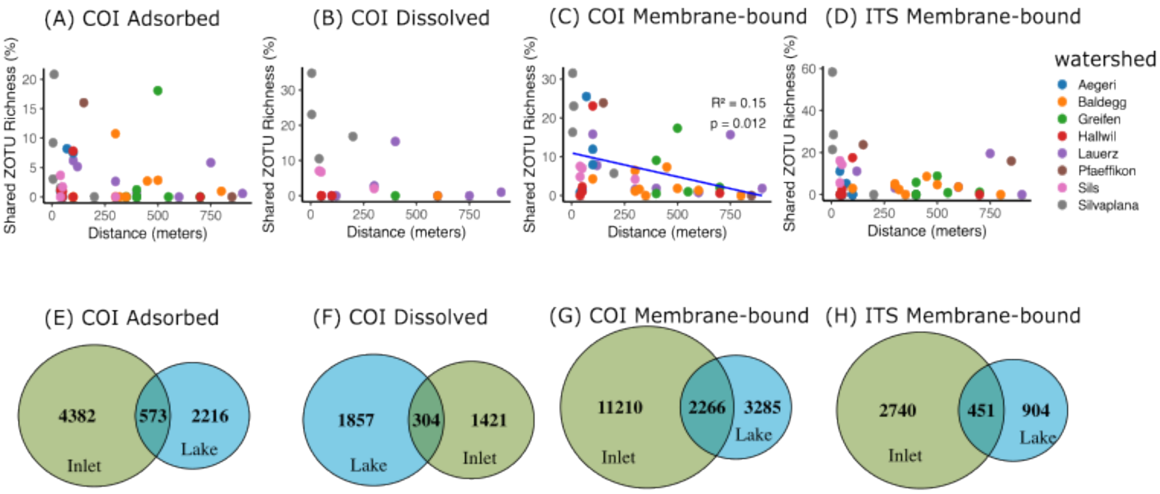


Figure S15: Percent shared richness of ASVs over distance between inlet and lake samples, color-coded by lake watershed for (A) Adsorbed, (B) Dissolved, and (C) Membrane-bound eDNA states for the COI marker and (D) Membrane-bound state for the ITS marker. Blue line: linear regression with $R^2$ and p values shown. The absence of a trend line indicates no significant relationship. Venn diagrams show unique and shared ASVs between lake and intlet samples for (E) Adsorbed, (F) Dissolved, and (G) Membrane-bound eDNA states for the COI marker and (H) Membrane-bound state for the ITS marker.

Table S6: Kruskal-Wallis test parameters of observed ASV richness to see a significant difference between stream, inlet, lake, and outflow samples. Follow-up Dunn's post hoc test results are shown in Table S7. This data is visualized in Figure 6.

| Marker | State | n | df | chi-squared | p |
|---|---|---|---|---|---|
| COI | Adsorbed | 218 | 3 | 7.34 | 0.062 |
| COI | Dissolced | 148 | 3 | 20.11 | 1.61E-4 |
| COI | Membrane-bound | 215 | 3 | 60.34 | 4.30E-13 |
| ITS | Membrane-bound | 221 | 3 | 10.51 | 0.015 |

Table S7: Dunn's post hoc test parameters showing significant differences in ASV richness between stream, inlet, lake, and outflow samples conducted as a follow-up to the Kruskal-Wallis test (Table S6).

| Marker | Panel | Comparison A | Comparison B | Raw p | Adjusted p | Significance |
|---|---|---|---|---|---|---|
| COI | DIS | up-stream | Inlet | 0.771 | 0.771 | ns |
| COI | DIS | up-stream | Lake | 0.00001 | 0.00008 | **** |
| COI | DIS | up-stream | outflow | 0.331 | 0.497 | ns |

| | | | | | | |
|---|---|---|---|---|---|---|
| COI | DIS | Inlet | Lake | 0.00141 | 0.00423 | ** |
| COI | DIS | Inlet | outflow | 0.453 | 0.543 | ns |
| COI | DIS | Lake | outflow | 0.232 | 0.463 | ns |
| COI | MEM | up-stream | Inlet | 0.466 | 0.559 | ns |
| COI | MEM | up-stream | Lake | 0 | 0 | **** |
| COI | MEM | up-stream | outflow | 0.00391 | 0.00587 | ** |
| COI | MEM | Inlet | Lake | 0 | 0 | **** |
| COI | MEM | Inlet | outflow | 0.00205 | 0.0041 | ** |
| COI | MEM | Lake | outflow | 0.697 | 0.697 | ns |
| COI | MEM | up-stream | Inlet | 0.506 | 0.607 | ns |
| ITS | MEM | up-stream | Lake | 0.00331 | 0.0198 | * |
| ITS | MEM | up-stream | outflow | 0.0873 | 0.175 | ns |
| ITS | MEM | Inlet | Lake | 0.0582 | 0.175 | ns |
| ITS | MEM | Inlet | outflow | 0.184 | 0.276 | ns |
| ITS | MEM | Lake | outflow | 0.771 | 0.771 | ns |

Table S8: Results of environmental vector fitting (envfit) showing the strength of association ($R^2$) and significance (p-value) of environmental variables with NMDS ordinations for each DNA state and marker.

| Marker | State | variable | r2 | p_value |
|---|---|---|---|---|
| COI | ADS | temp | 0.28469191 | 0.001 |
| COI | ADS | EC | 0.07503004 | 0.001 |
| COI | ADS | ppm | 0.07482953 | 0.001 |
| COI | ADS | pH | 0.05300151 | 0.001 |
| COI | DIS | temp | 0.16694651 | 0.001 |
| COI | DIS | ppm | 0.0938909 | 0.001 |
| COI | DIS | EC | 0.08842709 | 0.001 |
| COI | DIS | pH | 0.04272215 | 0.001 |
| COI | MEM | temp | 0.50296538 | 0.001 |
| COI | MEM | ppm | 0.26475728 | 0.001 |
| COI | MEM | EC | 0.2611332 | 0.001 |
| COI | MEM | pH | 0.0554898 | 0.001 |
| ITS | MEM | temp | 0.14558052 | 0.001 |
| ITS | MEM | pH | 0.04376525 | 0.001 |
| ITS | MEM | ppm | 0.01617898 | 0.001 |
| ITS | MEM | EC | 0.01523819 | 0.001 |
